# Structural basis for UDG identifying DNA damage in the nucleosome

**DOI:** 10.1101/2025.04.17.649289

**Authors:** Safwen Ghediri, Ralf Blossey, Fabrizio Cleri

**Affiliations:** Institut d’Electronique Microelectronique et Nanotechnologie (IEMN CNRS UMR8520) and Département de Physique, Université de Lille, 59652 Villeneuve d’Ascq, France; Unité de Glycobiologie Structurelle et Fonctionnelle (UGSF CNRS UMR8576), Université de Lille, 59000 Lille, France; Laboratory for Integrated Micro Mechatronics (LIMMS CNRS IRL2820) and University of Tokyo, Komaba, Meguro-Ku, Tokyo 153-8505, Japan

## Abstract

The DNA base-excision repair (BER) pathway is initiated by a glycosylase, such as UDG, which identifies and removes a wrong base incorporated in the DNA sequence. The very early steps in the identification of the DNA damage are crucial to the correct initiation of the repair chain, and become even more complex when considering the realistic environment of damage to the DNA in the nucleosome. We performed all-atom docking and molecular dynamics computer simulations of the interaction between the glycosylase UDG and a mutated uracil. The model system is a whole nucleosome in which DNA damage is inserted at various positions along the 145-bp sequence. It is shown that damage recognition by UDG requires very strict structural conditions, unlikely to be matched by purely random search along the DNA. We propose that mechanical deformation of the DNA around the defective sites may help signaling the presence of the defect, accelerating the search process.

## 1 Introduction

The base-excision repair (BER) pathway typically requires four or five enzymes to perform the DNA repair process. It is initiated by removing the damaged base by one of the eleven known mono- or bifunctional glycosylases [1], depending upon the type of defect. This process forms apurinic or apyrimidinic (AP) sites. For example, the RNA base uracil may appear in DNA through misincorporation by DNA polymerase, giving A:U pairs, or via cytosine deamination, which creates promutagenic G:U wobble mispairs. Uracil-DNA-glycosylase (UDG) cleaves the uracil base from both A:U pairs and G:U mispairs thereby creating an AP-site. Such hanging AP-site is then cleaved by an AP-endonuclease enzyme, such as APE1, which generates 3′-OH and 5′-dRP termini at the break site. The subsequent step of filling the single-nucleotide gap can follow either a long-patch, or short-patch repair path. The latter is generally considered to be the dominant choice, since it requires replacing a single nucleotide by DNA polymerase-*b*, therefore being faster and less prone to errors. On the other hand, long-patch repair is observed in post-replicative BER initiated by UNG2 or NEIL1 glycosylases, typically expressed during the S-phase [2]. Long-patch repair is also observed when the 5′-dRP terminus is oxidized to different abasic lesions (e.g., dioxibutane [3], C4′- or C2′-AP [4]), in which cases Pol-*b* can excise the damaged terminus with reduced efficiency.

As it clearly appears, the different steps of any DNA repair chain depend on a number of microscopic details, which are essential in determining the successful outcome of the entire process. Even a few “wrong” atoms in the “wrong” place can stop the process of DNA healing, with drastic consequences on the cell survival. For example, a 3′-dRP (deoxyribose phosphate) termination at the break site is very efficiently cured by Pol-*b*, whereas tyrosyl-DNA phosphodiesterase TDP1 can slow its action from less than a minute down to several hours, if the damaged 3′-DNA end is instead terminated by a phosphoglycolate [5] (as it often happens after radiation damage in presence of O_2_), despite the dRP looks a more chemically complex species than PPG.

The atomistic details of the interactions can be revealed by experimental studies of protein-DNA interaction at the scale of the single molecule, such as x-ray diffraction, nuclear-magnetic resonance, electron cryo-microscopy. These are quite difficult as well as resource-intensive techniques, and typically involve the active fragment of the (often quite big) test protein and a very short target DNA oligomer. By contrast, DNA in the cell nucleus is typically packaged as “beads-on-a-string” into nucleosomes, the building blocks of chromatin. About the histone octamer, a stretch of ∼50 nm of DNA (145 to 147 base-pairs) is continuously wound to form a nucleosome core particle (NCP) of about 11 nm diameter. In turn, pairs of nucleosomes are connected by stretches of “linker” DNA, with length variable between some 20 to about 200 base-pairs (that is, about ∼5 to ∼50 nm). Therefore, the free-DNA short oligomers typically used in experiments can only be a poor representation of the actual DNA in the chromatin; even the long DNA linkers could not be represented as free floating fragments, but are entropically and mechanically constrained by their ends attached to each pair of nucleosomes. Moreover, since on average about 75% of the DNA is wrapped into nucleosomes, the external damage such as radiation- or chemically-induced should be statistically more often localized at nucleosomes. Given such a context, molecular computer simulation could be of great help to shed light on the detailed mechanisms of action of the many physical-chemical steps involved in DNA repair in such a complex supramolecular structure. In this work we explore by means of combined docking and molecular dynamics (MD) computer simulations, the DNA-UDG interactions in the full environment of an isolated nucleosome (actually a NCP, i.e., a complete nucleosome core without linkers). We manually introduce constructs of DNA damage at various sites around the nucleosome, in the form of a uracil base. One basic question is underlying, namely, whether the enzyme actively modifies the defect site during the recognition, or rather the defect site “exposes” itself to facilitate the identification. For UDG to target the uracil, the wrong base must be flipped out of the DNA axis in extrahelical position, such that the UDG binding pocket can access it and proceed to the excision. However, this same question concerns in general any enzyme searching for some active site on the DNA. Depending on the ordering of the elementary steps of the process, such an interaction could represent either a “bind–then–bend” (the enzyme actively deforms the defect site after finding it), or rather a “bend–then–bind” (the defect is spontaneously modified prior to enzyme binding) sequence [6–11]. It may be expected that this issue becomes further complicated when DNA is wrapped around the nucleosome, because of the steric hindrance and extra charges provided by the histone proteins, and notably the highly flexible histone tails. As a matter of fact, very few experiments are available to date for UDG interacting with a nucleosome, e.g. [12–16], none of which provided detailed molecular structures of the full protein-nucleosome complex. This underscores the importance and opportunity of addressing these issues also by computer simulation, at least to provide a first glance about the numerous open questions.

We already addressed in two recent works the computational study of this same enzyme-DNA complex. We started [17] by first looking at the detailed energetics and kinetics of extrahelical uracil flipping in a short DNA oligomer, by performing free-energy calculations with a metadynamics approach; then, some preliminary results were presented in a comparison between UDG and PARP1, as representative of the BER/SSBR partly-shared repair pathway [18]. With the present work, we want to provide a thorough investigation of UDG-DNA interaction in the realistic environment of a whole nucleosome, possibly more relevant to the development of DNA damage repair within the nuclear chromatin.

## 2 Methods

To elucidate the molecular interactions in a model nucleosome between UDG and a flipped-out uracil, we designed a complete simulation protocol, starting with (i) the design of the atomic structure of the nucleosome, DNA and protein fragments; then (ii) docking of the rigid fragments (protein to the DNA) with an approximate free-energy functional; followed by (iii) all-atom molecular dynamics (MD) simulations with a state-of-the-art empirical force field of the complete protein+nucleosome, embedded in a large box of water and ions for a physiological solution, and (iv) detailed analysis of the molecular configurations and structural dynamics.

### 2.1 Docking

Docking of proteins and nucleic acids is still a rapidly evolving field of research. We tested different methods available as online server platforms: HADDOCK [19, 20], HDOCK [21] and the very recent pyDockDNA [22]. For the UDG/DNA interaction, after testing different methods among the available online platforms, pyDockDNA turned out to be the only one capable of providing reliable results. We started the docking using the molecular structure of UDG from RCSB Protein Database entry **1EMH**, and a DNA-X generic fragment of seven base-pairs including a flipped uracil or thymine, from our previous study; for each of the six interaction sites along the nucleosome (see below), we replaced the DNA-X sequence with the local 7-mer sequence, however without allowing for structural relaxation. To enhance accuracy, we incorporated the position of the UDG pocket interacting with the flipped nucleotide into the docking parameters. pyDockDNA generated the top 100 docking results, from which we selected the most suitable candidate, based on a comparison (minimum RMSD) of the flipped-nt geometric positioning in the pocket with respect to the experimental structures **1SSP** and **1EMH**. Then, the selected UDG+7mer complex was repositioned back in its nucleosome site, to start the MD equilibration.

### 2.2 Molecular dynamics

All the MD simulations and most data analysis, were performed with the GROMACS 2020.4 package [23, 24]. The Amber14 force-field was used in the simulations [25]; notably, Amber14 already includes the latest Parmbsc1 extension for nucleic acids [26]. To ensure the primary stability of the post-docked configurations, we first performed energy-minimization using a steepest-descent algorithm, followed by {NVT}-{NPT} equilibration for 150 ps, to bring the systems at 310K and 1 bar pressure. The ensembles of DNA-proteins were solvated in water boxes of typical size 13x13x13 nm^3^, with about 75,000 water molecules, along with Na^+^ and Cl^-^ ions for neutralizing the system charge and maintaining a physiological 0.1M salt concentration. In some instances we also performed rapid annealing and quenching cycles, to improve the mutual positions of protein and DNA, for example after a manual adjustment necessary to remove a steric clash. Each cycle was typically performed by ramping up the temperature from 310K to 400K, and then back to 310K, in steps of 10K for 500 ps each. Usually, during the annealing cycle the DNA was frozen in its conformation and only the protein was left free to adjust.

Coulomb forces were summed with particle-mesh Ewald sum (PME), using a real-space cutoff of 1.2 nm, equal to the cut-off radius of shifted Van der Waals potentials. We used rigid bonds for the water molecules, which allowed integration of the equations of motion with a time step of 2 fs for the thermal equilibration phases, and 1 or 2 fs for production runs.

Trajectory clustering analysis was performed using a special subprogram of GROMACS. The MD simulation stores ’frames’ containing the positions, velocities and forces of all particles in the system, at prescribed intervals (typically every 10 to 50 ps, or longer: a 1 *µ*s-long MD run can store as much as 100,000 frames of about 2Mbytes each, resulting in data files with size of hundreds of Gbytes); the subprogram calculates a matrix of root-mean-squared displacements (RMSD) between each pair of frames, by comparing the positions of each atom in the pair; then, RMSD values are grouped according to a cut-off criterion, and clusters of similar frames, typically separated by a small enough RMSD, are detected. The cluster(s) with the highest number of members, and those which are more frequently sampled over the entire simulation time, provide an indication of the ’best-average’ molecular configurations, thus allowing to bypass the noise of the short-time atomic fluctuations.

Free-energy analysis and contact surface calculations were estimated by the PDBePisa web-based utility [27], by extracting and averaging a few selected configurations in PDB format from the MD trajectory. The resulting values are approximated, however they can provide at least a qualitative appraisal of the binding energies and chemical affinity involved in the protein-DNA interaction.

Metadynamics simulations of extra-helical nucleotide flipping were performed with the PLUMED plugin to GROMACS [28], according to the protocol described in our previous work [17]. The principal collective variable (CV) to force the extrahelical base flipping was defined by the dihedral connecting the four points *P*_*i*_ representing the centers of mass of, respectively, the four adjacent and facing bases (*P*_1_), the two phosphate groups of the adjacent nucleotides (*P*_2_ and *P*_3_), and the heavy atoms of the flipping nucleotide (*P*_4_); *P*_1_, *P*_2_, *P*_3_ form one triangle and *P*_2_, *P*_3_, *P*_4_ a second triangle sharing the *P*_2_ - *P*_3_ side (see also [29]); such an arrangement allows the point *P*_4_ (and the whole central nucleotide) to rotate between ±180° about the phosphate backbone. In the course of the study, also a second CV was defined, to bring the uracil as close as possible to the UDG active site (see below).

### 2.3 Nucleosome

The initial molecular configuration of the nucleosome is taken from the RSCB Protein Database, entry **7OHC** [30]. This is a cryo-microscopy structure of the entire histone octamer with 145 DNA base-pairs resolved at an average RMS of 2.5 Å, reconstituted from human nuclear extract expressed in *E. coli*; the missing histone residues were completed with the Modeller utility of SwissProt [31] to obtain a complete nucleosome including the full-length histone tails. The 145-bp DNA has the artificial Widom 601-sequence [32], originally selected to maximize the binding affinity to the histone octamer; to obtain a model structure useful for our computer simulations, we removed all the crystallization water molecules. DNA in the nucleosome is wrapped left-handed about the histone core, making two nearly complete turns (‘gyres’) that join at the dyad symmetry point; the two DNA turns define two circles lying in two ideally parallel planes, with a superhelical symmetry axis (SHL) perpendicular to the center of the circles; the relaxed DNA double helix makes a complete twist around its double-helical axis, about every 10.4 bp; correspondingly, 14 contact points between DNA and the histone core proteins can be identified within the nucleosome structure, loosely situated at the minor groove locations facing inwards. Based on these standard geometrical features, we defined the sites where to place DNA damage in the form of either a uracil (oxydised cytosine), or a single-strand break (SSB), in structurally significant positions (see discussion below). Figure 1 provides a summary of the nucleosome structure and uracil locations.

**Figure 1.**
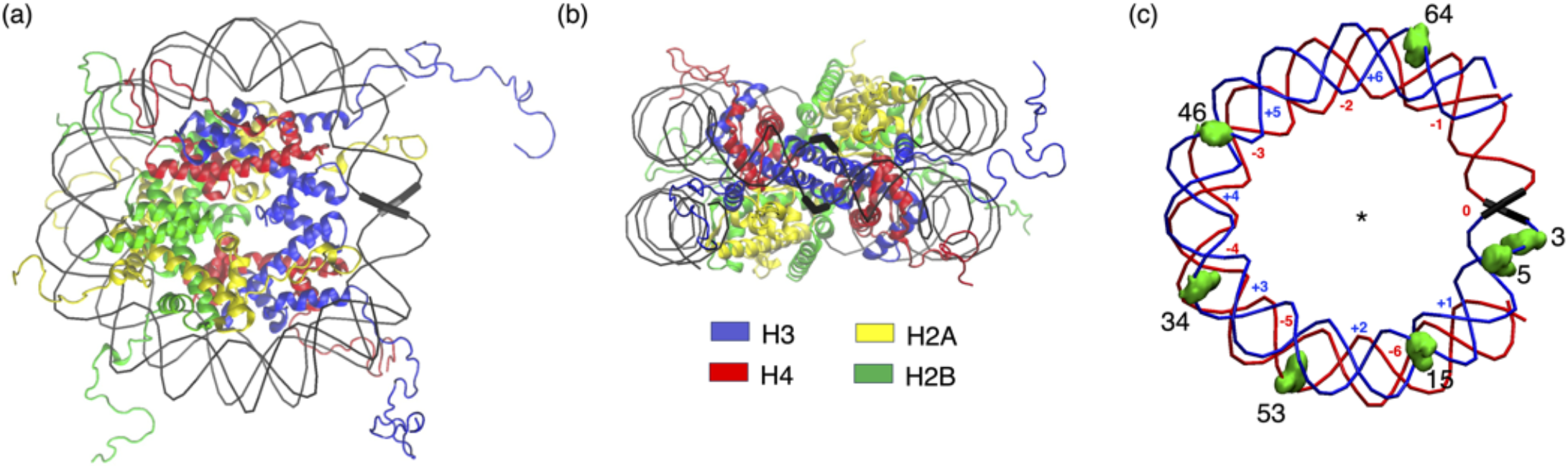
**(a)** Top view of the model nucleosome 7OHC completed with histone tails, after 5-ns molecular dynamics equilibration at T=310K; histone proteins represented as color ribbons; DNA as a simple black wire; the dyad is indicated by a thicker black wire. **(b)** Side view from the dyad, showing the antisymmetric arrangement of the four histone dimers. **(c)** UDG/DNA interaction sites, with the positions of the uracil substitution indicated by green spheres, and numbered according to their location along each nucleotide chain (see also Table 1); the two axisymmetric DNA double-strands start from the dyad (nucleotides 0) and proceed clockwise “below” (red), and counterclockwise “above” (blue) the central, mirror-symmetry plane; the superhelical locations (minor-groove inner contacts) are labelled in red and blue, correspondingly; the central asterisk indicates the position of the superhelical axis, perpendicular to the page.

## 3 Results

### 3.1 Structures of UDG and flipped-out nucleotides

To obtain a fully-formed interrogation complex (see also below, sect.3.4), the uracil defective nucleotide is initially arranged in a fully-extrahelical position, based on the findings of our metadynamics study on short DNA oligomers [17], i.e. rotated at 180° about the O-P-O phosphate chain with respect to its equilibrium position. This conformation could equally represent the uracil, either spontaneously fluctuating before the interaction with UDG (in a “bend–then–bind” sequence [6, 7]), or right after UDG performed a pinch–and–pull mechanical action (as in a “bind–then–bend” sequence [33, 34]). To assess the physical-chemical selectivity of UDG towards the mutated vs. non-mutated sequence, we also ran MD simulations on identical configurations, by replacing uracil with thymine or cytosine.

**Table 1.**
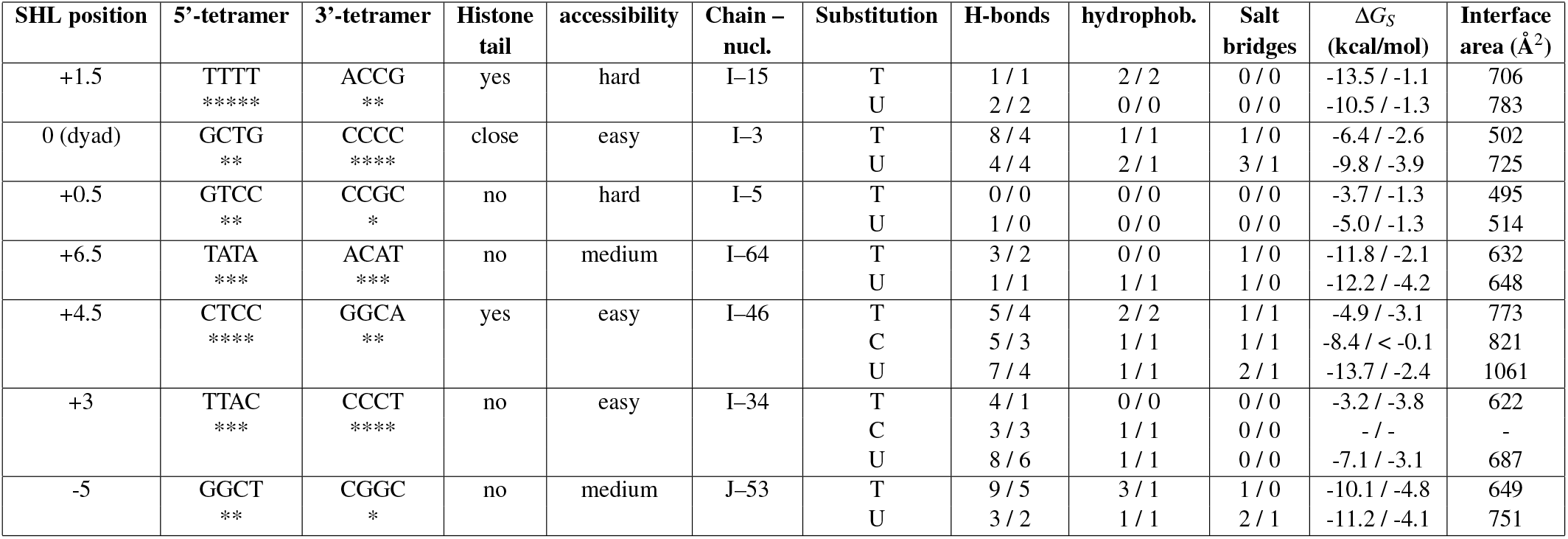
Summary of the molecular interactions of UDG with each flipped-out nucleotide (T/C vs. U), for the various positions around the nucleosome, averaged over the last 10 ns of each 100-ns MD trajectory. For H-bonds, salt bridges, hydrophobic interactions, and D*G*_*S*_, the first number is the total and the second that corresponding to the flipped nucleotide only. The number of ’*’ under the flanking tetramer sequence gives a (rough) estimate of the local flexibility of the ssDNA strand.

For the molecular structure of the human UDG glycosylase, we started from two experimental structures of the enzyme interacting with a short DNA oligomer, available in the RCSB PDB archive, the **1EMH** and the **1SSP**; the two are very similar, in that they both include a wild-type UDG bound to a DNA decamer containing in the 5th base, respectively, an uncleaved uracil [35], or a cleaved uracil [36]. It is worth noting that the two molecular structures are nearly identical, with the cleaved U displaced by just 0.12 nm inside the UDG pocket after the cut.

As said, a set distinct configurations of the nucleosome structure were generated, each featuring two(three) variations: one with thymine (T) and the other with uracil (U); in some cases we also tested a cytosine (C) variant. The substitution of T/C by U was performed using the Chimera software [37]. The selection of specific positions for these modifications was guided by geometrical criteria. The level of accessibility was defined based on two factors: (i) the solvent exposure of the nucleotide, i.e., whether it is oriented toward the interior of the nucleosome (hard), thereby restricting UDG access; or facing outwards (easy), where it remains fully exposed; or intermediate between the two (medium); and (ii) the eventual proximity of a histone tail, that may perturb UDG while interacting with the flipped nucleotide. This criterion allowed for a systematic evaluation of how nucleosomal positioning influences the recognition and excision of uracil by UDG.

In principle, uracil in U:A base-pairs within straight dsDNA present the same pattern of hydrogen bond donors and acceptors as T:A basepairs (in fact, DNA polymerase cannot discriminate between the two [38]), but differ by lacking the thymine 5-methyl group. Uracils in U:G mispairs, however, are shifted towards the major groove (“wobble”) and leave partial hydrophobic gaps within the DNA base stack. In addition to these structural features, UDG has been supposed to induce a structural distortion on the DNA, to further distinguish uracil-containing nucleotides from bulk DNA. Although these interactions are transient and likely difficult to stabilize for experimental studies, crystal structures of UDG alone, in complex with DNA, and in complex with the glycosylase inhibitor Ugi (see e.g. [39]), coupled to biochemical and biophysical data have revealed clues to this critical interaction.

### 3.2 Nucleosome accessibility and UDG positioning

Compared to the ’clean’ experiments carried out with short DNA oligomers, the nucleosome presents UDG with a much more rugged landscape: the DNA is bent with a tight curvature, the major/minor groove widths, base-pair and stacking parameters are fluctuating to rather different values from those of the ideal, straight B-DNA. Confronting the set of different interaction sites between UDG and a flipped nucleotide along the nucleosome, allows to understand at least some necessary conditions for a correct binding and subsequent excision action of the enzyme. (Here “correct” means an arrangement as much comparable as possible to the experiments, e.g. [35, 36, 40], which depict the final (“Michaelis” [41]) state of the DNA-enzyme interaction.)

The initial docking chiefly aimed at positioning the flipped nucleotide in the correct position inside the binding pocket. However, this microscopic configuration can be achieved with many different macroscopic arrangements of the whole UDG about the DNA. One evident structural feature of UDG is its central, wide beta-sheet motif, formed by four parallel beta-strands lying in a common plane also including a short alpha-helix (residues 122-132), which makes up one side of the binding pocket (Figure 2c). In all cases in which a correct interaction is observed, this motif (for example, identified by the line *D* joining the *a*-carbon of residues 126 and 264) lies off the perpendicular to the central axis of the interacting DNA strand, with a slight inclination at about 70-75° (see Figure 2a). A second, geometrical parameter characterizing the mutual UDG-DNA arrangement can be defined as the “angle of attack”: in the correct binding, the plane identified by the line *D* and, e.g., the central N3 atom of the flipped-out nucleotide, is also somewhat off the perpendicular to the DNA local main axis (see Figure 2b). However, these are just necessary but not sufficient conditions for the correct arrangement of UDG around the nucleosome. Such a correct arrangement allows at least to: (i) achieve a proper orientation of Leu272, which should enter the minor groove, very close to the phosphate of the flipped-out nucleotide, and sandwiched between its two neighbors 5′*/*3′ nucleotides; and (ii) to place the 146-149 and 211-216 loops of UDG facing the major groove, next to the flipped nucleotide. Notably, in order to preserve enough space for the central nucleotide to flip-out through the major groove, these two loops must keep a distance of at least 1.5 nm from the Leu272 loop (see dashed lines in Fig.2c). It will be seen that the geometrical constraints of the nucleosome structure do not always permit such ideal conformations of UDG about the DNA, thereby motivating the experimentally-observed rotational dependence of UDG catalytic activity in the nucleosome [12].

**Figure 2.**
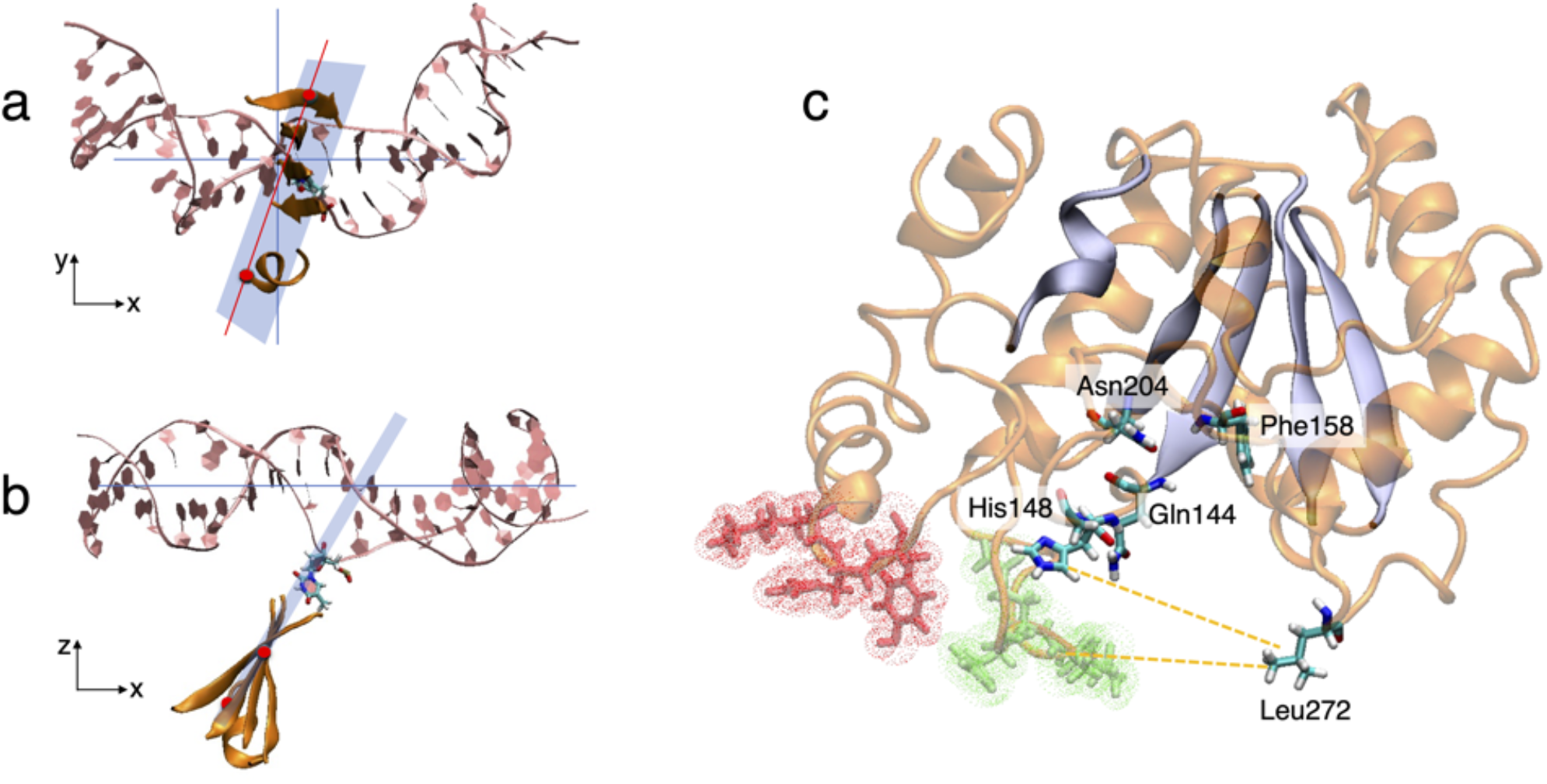
(**a**,**b)**) Two views [**(a)** rotated by 90 degrees about the horizontal x-axis, w/r to **(b)**] of the imaginary plane (blue) containing the beta-sheet motif of UDG (depicted in orange ribbons). **(a)** shows the inclination by about 70° of the *D* line (red) joining the two *a*-carbon atoms (red spheres) of residues 126 and 264. **(b)** shows the inclination of the blue plane also in the perpendicular direction. Note that the z-axis coincides with the SHL symmetry axis of the nucleosome. **(c)** The binding pocket of UDG. The beta-sheet and the *a*-helix 126-132 are highlighted in blue; Asn204 and Phe158 are the a.a. providing the key interaction with the aromatic O4 of the flipped-out nucleotide inside the pocket; Gln144 is the a.a in charge of nucleophilic attack to the glycosidic bond on the flipped nucleotide; Leu272 is the minor-groove intercalating a.a.; His148 is the central a.a of the 146-149 loop that broadly interacts with the DNA major groove; the other major groove-facing loop 212-215 is highlighted in green; the side loop 114-116 is highlighted in red. The orange-dashed lines indicate the “opening” of the binding pocket, as the average distance between Leu272 and the major-groove protein loops.

Coming to the physical chemistry of the interaction, the flipped-out nucleotide must fit inside the binding pocket, right next to the main beta-sheet, where it should get bound by H-bonds with the two key residues Asn204 and Phe158. If the base is uracil, then Gln144 helped by His268 will make the nucleophilic attack that cuts the glycosidic bond; otherwise the base will be ignored. (In practice, UDG may also excise a correct pyrimidine, however the rate of such errors is in the range 10^−8^ [42]). Note that given the size of the pocket, purines cannot fit in. As it will be seen, although the initially-docked complexes may generally display a rather correct positioning of the flipped nucleotide in the binding pocket, the subsequent dynamics shows that the interaction can only be maintained when the geometrical constraints on UDG-DNA steric arrangement are satisfied.

Notably, the UDG positioning can also display a variable degree of sequence-dependence, since the extrahelical flipping induces a substantial phosphate backbone deformation, at least of the base-pairs immediately adjacent the 5′ and 3′ sides, the DNA getting kinked at the flipped site. On very general grounds, AA/TT dimers in the DNA sequence tend to favor intrinsic bending, whereas GC dimers are generally more rigid and resist bending. At a slightly longer distance, e.g., adjacent tetramers, the sequence-dependent stacking interactions influence the overall DNA persistence length and flexibility: from the most flexible to the most rigid, we have Poly(T) > Poly(C) > Poly(A) ∼ TATA > GC-rich sequences > Poly(G) (G-quadruplex) (see Ref. [43] and references therein). Our chosen interaction sites along the nucleosome have quite different flanking tetramer sequences; Table 1 below provides an estimate of the corresponding flexibility, defined as average of stretching, bending and twisting constants [44]. Given the important deformations of the DNA backbone observed, such different elastic properties should impact on the relative ease of UDG positioning during its search for defects along the nucleosome.

Table 1 summarizes a number of results for the different UDG-DNA interaction sites calculated with the PDBePisa utility [27], by averaging over the last 10 ns of each MD trajectory. Each row corresponds to a substitution in the DNA chain/nucleotide indicated in column 6 (for example, the first row is a flipped-out thymine in chain I, position 16 (corresponding to SHL+1.5), the second row is a uracil in the same position, and so on). Whenever two values are shown separated by a “/”, the first refers to the total, and the second to the interaction with only the flipped nucleotide. D*G*_*S*_ in column 11 indicates the solvation free-energy gain upon formation of the interface, in kcal/mole (not a *total* free-energy of the compound formation).

The value is calculated as difference in solvation energies between the isolated and interfacing structures: a negative D*G*_*S*_ corresponds to hydrophobic interfaces, or positive protein affinity. Note that this value does not include the effect of satisfied hydrogen bonds and salt bridges across the interface given in columns 8-10. The interface area in column 12 is calculated as the difference in total solvent-accessible surface areas (Å^2^), between the isolated and interfacing structures, and divided by 2.

In general, it may be noted that both the solvation energy and the interface area are systematically larger for the U vs. T/C substitutions. Also, looking at the fraction of H-bonds, the ratio to the total is always favorable to the U vs. T/C. However, by looking at the overall geometry after the 100-ns dynamics, we find that only the configurations SHL-0 (or I-3), SHL+3 (I-34) and SHL+4.5 (I-46) still display a correct, or near-correct binding. Note, however, that for the C substitutions all the interactions are with residues outside the UDG binding pocket.

### 3.3 Structural dynamics of UDG interactions in the nucleosome

#### Position SHL+3

Both T34 and U34 start from a quite correct UDG/DNA arrangement, Leu272 is intercalated in the minor groove between the flanking nucleotides 33-35, and H-bonds are formed between the flipped base and the Phe158 and Asn204. However, very soon in the dynamics the H-bonds are broken in the T34 (blue plots in Fig.3c), which eventually gets out of the binding pocket; at the same time, Leu272 is pushed out of the minor groove, and UDG makes a rigid rotation about its original arrangement on the DNA (Fig.3b). By contrast, U34 retains the correct binding and interaction configuration for the whole duration of the dynamical trajectory (see Fig.3a and red plots in 3c).

**Figure 3.**
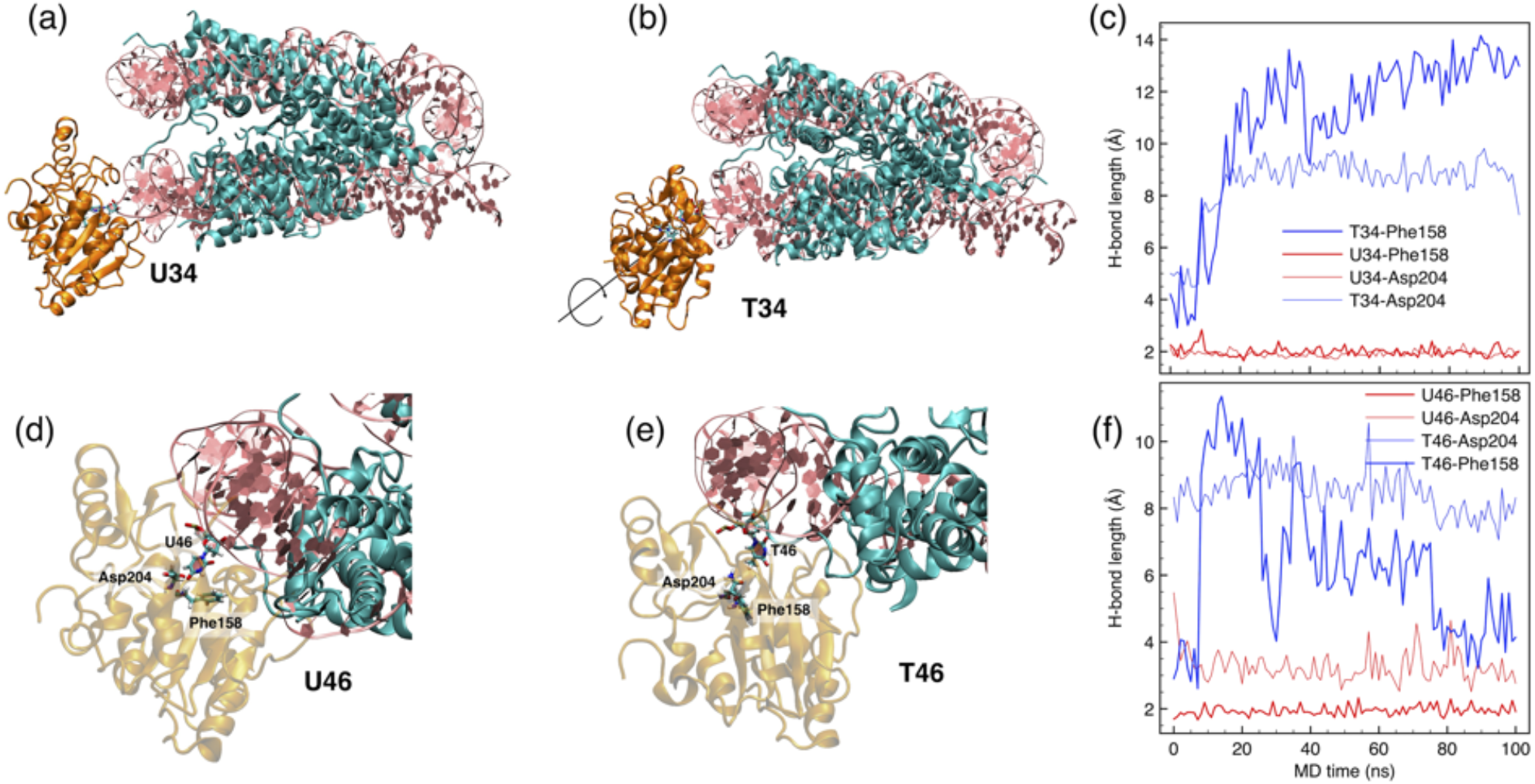
(**a**,**b)** Snapshots of UDG at position SHL+3, interacting with U34 and T34 nucleotides, after 100 ns of MD. DNA is depicted in pink ribbons, UDG in orange ribbons, histones in cyan ribbons. **(c)** Time evolution of the H-bonds initially established between the O4 atoms in the flipped out U/T (red/blue plots, respectively) with the N in Phe158 and N*d* in Asn2044. **(d**,**e)** Zoomed-in snapshots of UDG at position SHL+4.5, interacting with U46 and T46 nucleotides, after 100 ns of MD (same color-code). The flipped-out U and T nucleotides, and the a.a. Phe158 and Asn204, are highlighted as licorice sticks, colored according to element. **(f)** Time evolution of the H-bonds between O4 in U/T (red/blue) with N in Phe158 and N*d* in Asn204.

For this position we also tried the C34 substitution. It is worth noting that the terminal carbonyl C=O present in both U and T, is here replaced by the (pre-oxidation) amine NH_2_, rather bigger in steric size and electronic density. In fact, even if starting from a configuration with the flipped-out cytosine well inserted in the UDG binding pocket, and Leu272 correctly positioned in the minor groove, quite soon in the 100-ns MD simulation the C34 is pushed out of the pocket and starts flipping back into intra-helical conformation. Figure S1 in the supplementary material shows the position of C34 superposed at the beginning of the simulation (red) and after 100ns (blue); in the final stage of the dynamics, both His148 and Ala214 from the two UDG loops (red-green in Fig. 2c above) are receding, leaving enough room for C34 to flip back.

#### Position SHL+4.5

Also T46 and U46 start with comparable UDG/DNA arrangement and correct H-bonds; however, in this case Leu272 lies about the minor groove but away from the intercalation site, while in the major groove the loop 146-149 is closing the binding pocket. In such a configuration, it would be very difficult for the nucleotide U/T46 to flip-out, since it would sterically clash with the retracted position of His148. Such a shift of UDG with respect to the minor/major groove positioning is probably due to the interference of the N-terminal domain of histone H2A, which crosses here the minor groove and interferes with Leu272 insertion. Nevertheless, U46 maintains for the whole 100-ns its correct bonding inside the pocket with the key aminoacids Phe158 and Asp204 (Fig.3d), while T46 is rapidly pushed out of the pocket (Fig.3e) and loses contact (see H-bonds plots in Fig.3f). In the latter case, both the orientation angles (see Fig.2) are increased by ∼5-10 degrees, contributing to the loosening of the UDG/DNA contact.

Also for this position we tried the C46 substitution. The flipped-out cytosine starts again with a configuration well inserted in the UDG binding pocket; however, UDG is slightly displaced and Leu272 does not lie inside minor groove because of the already cited interference from H2A. In this case, during the 100-ns MD simulation the C46 is pushed out of the pocket but is unable to flip back into intra-helical conformation: even the small rearrangement of UDG about the nucleosome is already enough, for the two loops including His148 and Ala214 to block its way (see Figure S2 in the supplementary material). Given the clearly negative results obtained for the two positions SHL+3 and +4.5, we decided to not pursue further the C substitutions for the next nucleosome positions.

#### Positions SHL–5 and +6.5

These two rotational positions are rather similar, and indeed provide similar insight about a specific difficulty of UDG interacting with DNA in the nucleosome. Also in this case, the initial docked position of the flipped-out nucleotide in the binding pocket appears rather correct, for both the U/T pairs in positions 53 (SHL-5) and 64 (SHL+6.5). However, the problem is that after putting back the docked UDG+7mer at either position in the respective DNA turns, the side-loop 114-116 of UDG (see Fig.2c above) gets in contact with the DNA in the opposite nucleosome turn. As a representative example, Figure 4c displays a snapshot of the molecular structure for the U53 case, the others being all qualitatively similar. Notably, with UDG at SHL-5, the 114-116 loop (highlighted in red in the figure) makes contact with a Guanine-18 in SHL+3; and with a Guanine-12 in SHL-1.5, when UDG is at SHL+6.5. These contacts establish several dynamical H-bonds, which however make up a stable interaction on average.

**Figure 4.**
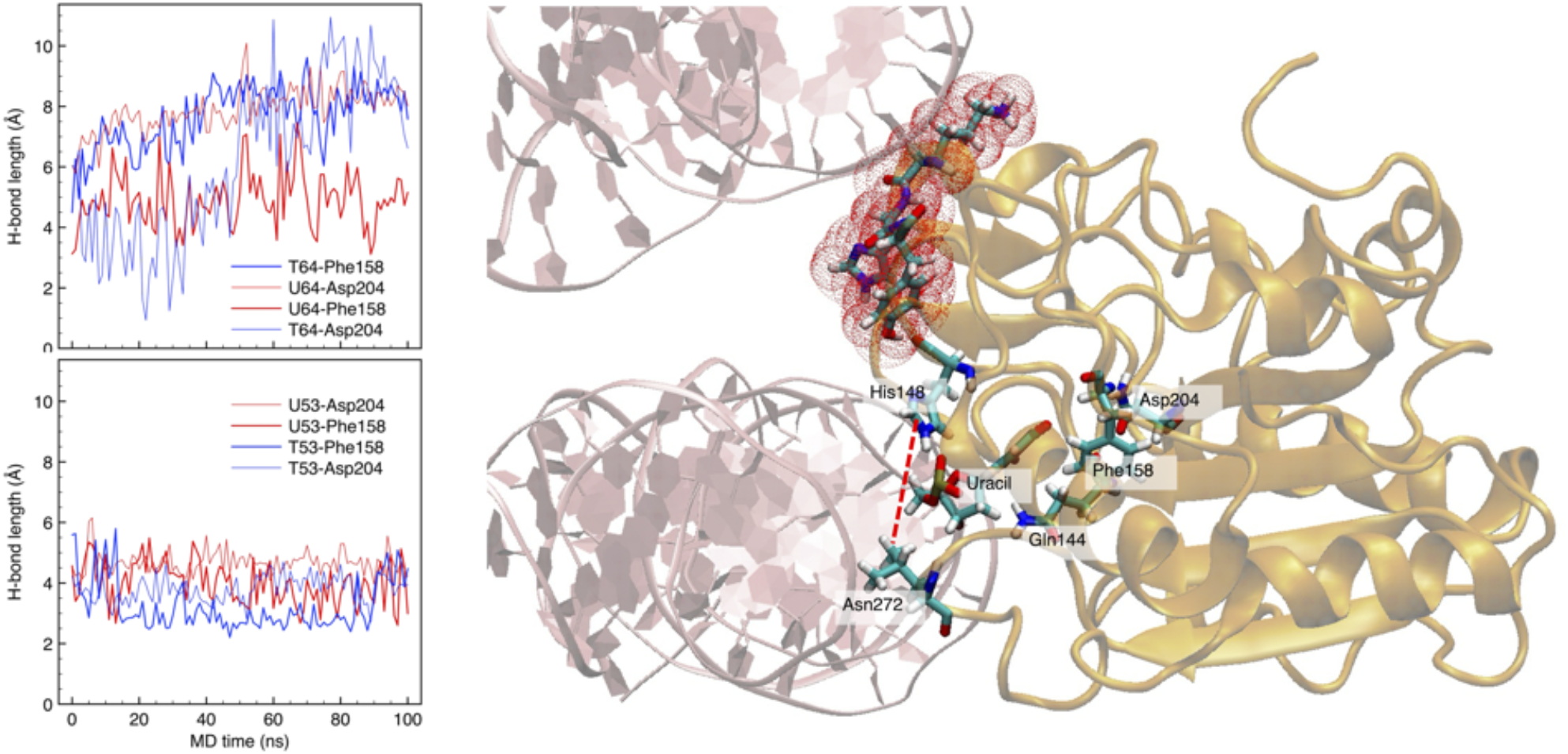
(**a**,**b)** Time evolution of the H-bonds between the O4 atom in the flipped out U/T (red/blue plots, respectively) with the N in Phe158 and N*d* in Asn204, at position SHL+6.5 U/T64 (**(a)** above), and SHL-5 U/T53 (**(b)** below). **(c)** Snapshots of UDG interacting with U53 after 100 ns of MD (same color code as in Fig.3). U53 is getting out of the binding pocket and lost contact with Phe158-Asp204. The side-loop 114-116 (highlighted in red, see also Fig.2 above) is in contact with the upper DNA turn and pushes the 146-149 loop deep in the major groove, thereby closing the pocket to an average width of 0.6-0.8 nm (see the red-dashed line joining His148-Leu272).

Such an arrangement forces the two UDG loops 146-149 and 212-215 (see again Fig.2c) deeper inside the major groove, while Leu272 remains rather correctly intercalated between the 5′*/*3′ flanking nucleotides. In this way, the access to the binding pocket is severely restricted, the opening width being on average well-below 1 nm, instead of the typical >1.5 nm (red-dashed line in Fig.4c).

As a result, both T and U remain inside the binding pocket, however the deformation imposed to UDG makes either of them to loose the correct alignment with the key a.a. (Gln144, Phe158, Asn204), as shown by the H-bond plots of Fig.4a,b for both the SHL-5 and SHL+6.5 positions. This would make the subsequent excision process pretty much ineffective. However, given the constraints, it is extremely unlikely that the central nucleotide (53 or 64) could even be flipped-out, in the first place, because of the numerous steric clashes it would face in its rotational movement across the major groove. It could still be possible that the flipping may occur via the minor groove, thus avoiding the clashes, however as we already demonstrated in our previous study [17], the corresponding energetic cost is much higher, the free-energy barrier increasing from D*G ∼*7 to 10 kcal/mol (that is, a Boltzmann probability exp(–D*G/RT*) decreasing by more than 100-fold).

#### Positions SHL-0 and +0.5

The dyad position is, at least in theory, an ‘easy’ accessible site: here only one DNA turn is winding about the histone core, and contact with histones is less close than in the rest of SHL positions. We tried two slightly different UDG interaction sites here: the arrangement of nucleotides U3/T3 nearly exactly coincides with the dyad site labelled “out” in the experimental study of Ref. [15], whereas the U5/T5 rather correspond to their “mid” dyad site. In that study it was reported that UDG at the outward facing site has a 15% efficiency (U-excisions/minute) compared to free DNA, and drops to 0.5% at the “mid” site, with moreover less than 10% total excision yield. Reduced enzyme efficiency, inversely proportional to the distance between the target site and the dyad axis, was traditionally explained with easier DNA accessibility provided by the partial, transient unwrapping at the nucleosome edges [45, 46]; however, we note that such hypothesis was built upon experiments on *isolated* nucleosomes, but such spontaneous unwrapping events would be infrequently observed in the dense chromatin structure.

For U3/T3 (facing outward) there is no direct contact between UDG and histones; however, the proximity to the DNA phosphate backbone of the H3 N-terminal tail and of a fold of the H4 histone, are already enough to perturb the correct arrangement of the enzyme in the interaction. Figure 5c shows the best configuration of U3 interacting with the binding pocket: it is observed that the two arginines from H3 and H4 are making contact with the DNA, at about SHL–0.5, the result being a constraint on the 146-149 loop of UDG, which pushes His148 to close the access to the binding pocket, the average opening as measured by the His148-Leu272 distance being < 1nm. As a result, the already flipped nucleotide can stay in place (see H-bond red plots in Fig.5a), however it would be hardly possible for it to flip by 180° into extra-helical, if starting from its regular intra-helical form. Besides being so strongly constrained, however, UDG retains a sharp discrimination ability between the uracil and the thymine base. Both the H-bond plots in Fig.5a, and the trademark interactions with Gln144 and His268 in Fig.5b that are the key step for the base excision, are clearly favorable to the uracil, while the thymine is practically pushed out of the binding pocket after the 100-ns dynamics.

**Figure 5.**
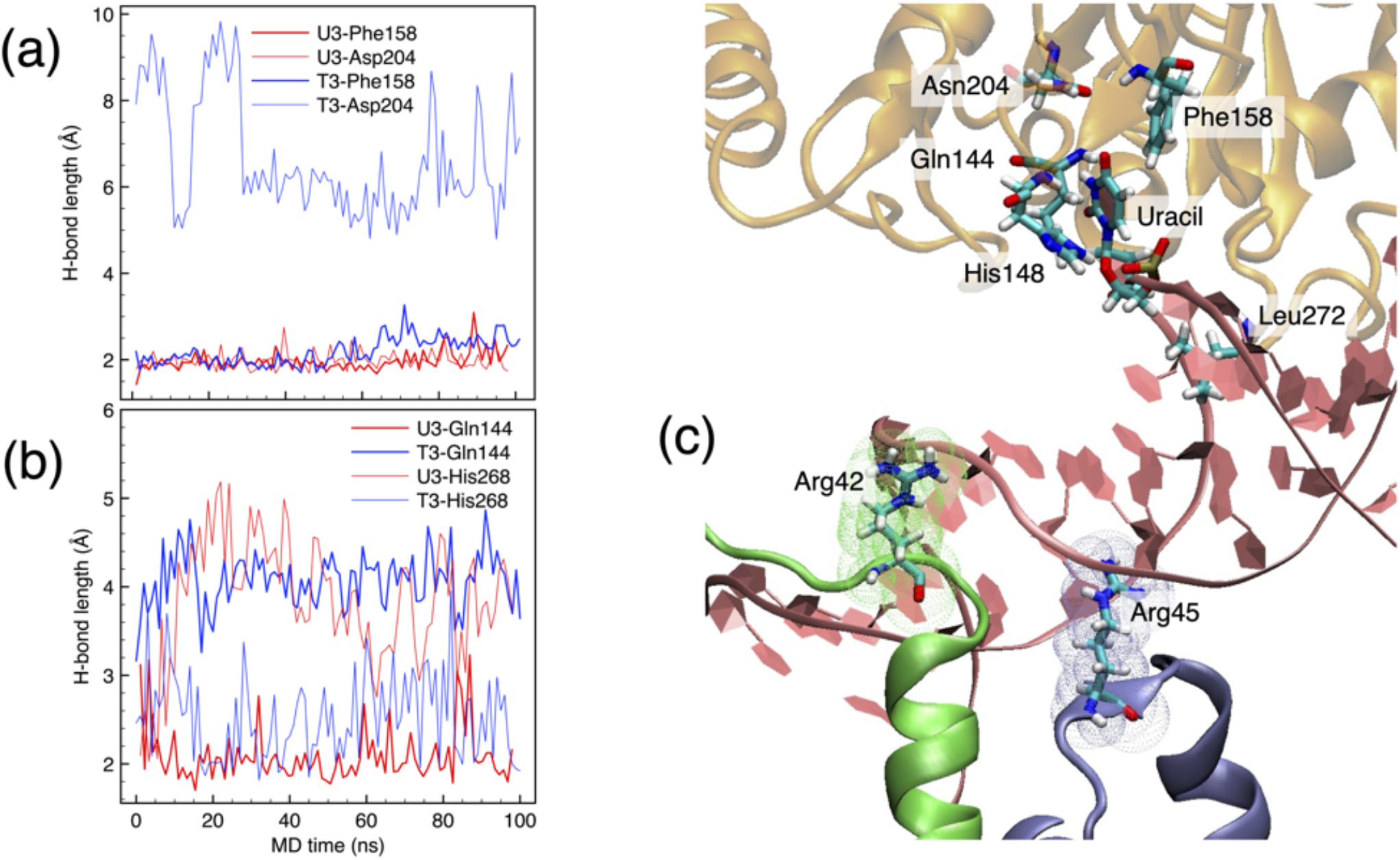
(**a**,**b)** Time evolution of the H-bonds between the O4 atom in the flipped out U/T (red/blue plots, respectively) with the N in Phe158 and N*d* in Asn204, at position SHL+6.5 U/T64 (**(a)** above), and SHL-5 U/T53 (**(b)** below). **(c)** Snapshots of UDG interacting with U53 after 100 ns of MD (same color code as in Fig.3). U53 is getting out of the binding pocket and lost contact with Phe158-Asp204. The side-loop 114-116 (highlighted in red, see also Fig.2 above) is in contact with the upper DNA turn and pushes the 146-149 loop deep in the major groove, thereby closing the pocket to an average width of 0.6-0.8 nm (see the red-dashed line joining His148-Leu272).

The close-by position U5/T5 (facing “above” the nucleosome) is right next to the histone H4 N-terminal tail (Figure 6a), whose contact with UDG forces a non-correct positioning at the DNA backbone. As a result, while Leu272 tries to keep the intercalation in the minor groove, the flipped-out nucleotide cannot properly accommodate in the binding pocket, as shown by the large distance maintained between the O4 atom of either U5 or T5, and the trademark H-bonding N and N*d* atoms of Phe158 and Asn204 (Fig.6b). Moreover, the flipped nucleotide is here flanked by quite rigid tetramers (Table 1), which contribute to a more difficult accommodation. Comparison of Fig.5a and Fig.6c clearly supports the experimental findings of Ref. [15], about the “out” vs. “mid” excision efficiency and total yield.

**Figure 6.**
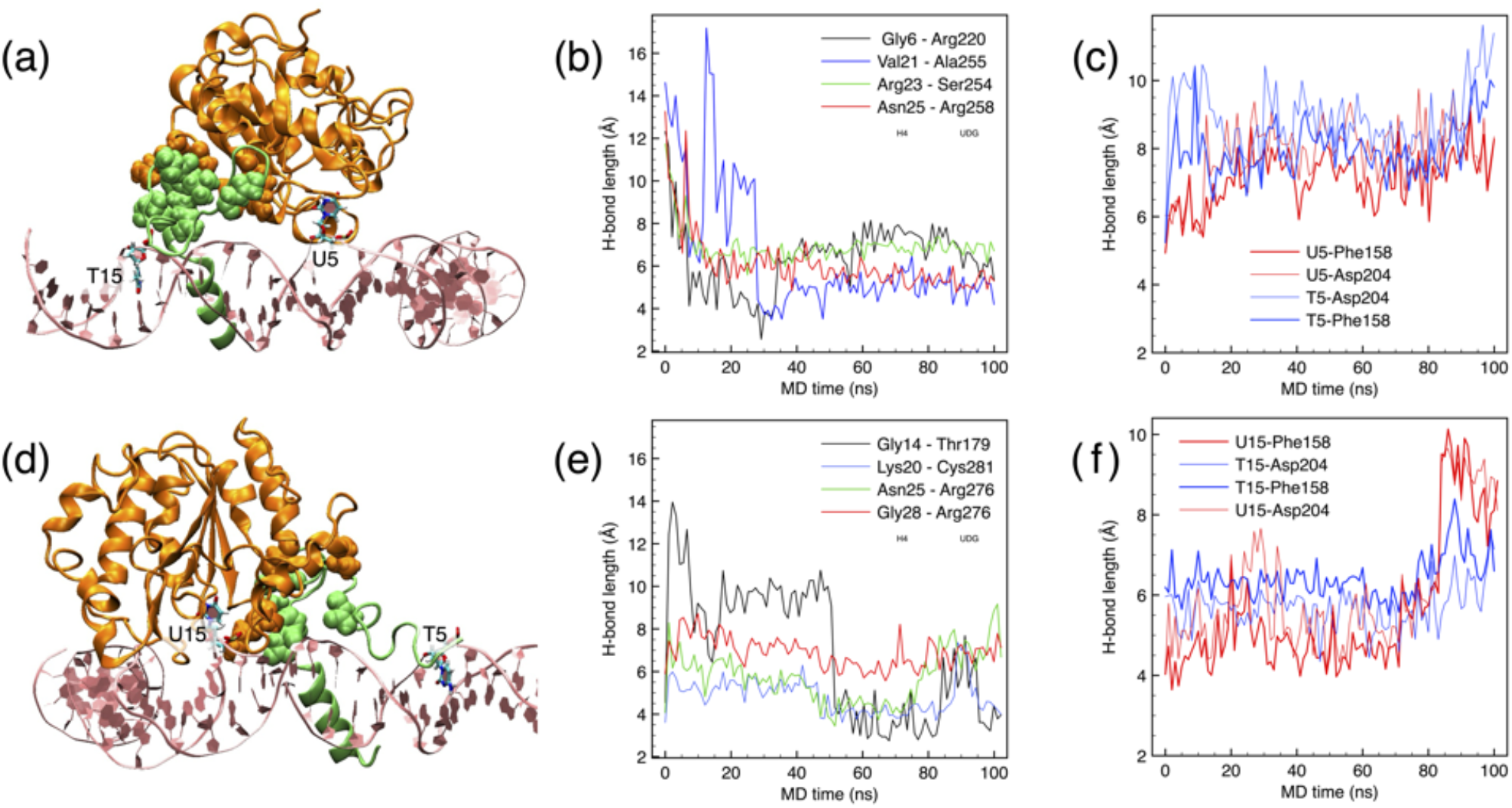
**(a)** Best cluster for UDG at uracil in position U5 (SHL+0.5). UDG in orange ribbons and histone H4 in green ribbons; the contacts between UDG and H4 are highlighted by VdW spheres; the site T15 is also shown for reference. **(b)** Time evolution of the contacts, represented by the distance between the centers of mass of the respective amino acids. **(c)** H-bonds between the O4 atom in the flipped out U/T (red/blue plots, respectively) with the N in Phe158 and N*δ* in Asn204. **(d-f)** Same as (a-c) for UDG at uracil in position U15 (SHL+1.5), same color codes.

#### Position SHL+1.5

This is another example of difficult positions for fitting UDG (“hard” in Table 1 above), the flipped nucleotide being at the last, barely accessible position next to the histone core; moreover, the

N-terminal of histone H4 runs through the minor groove close to the target DNA site (see Fig.6d). Being one full double-helix turn away from the previous SHL+0.5, UDG is now found on the opposite side of the same H4 histone (compare 6a and 6d). The histone tail interacts with the side chains of UDG (see Fig.6e), forcing a non-ideal conformation of the enzyme. Also in this case, like for the SHL+0.5 above, the distance between the flipped nucleotide and the key residues 158/204 in the binding pocket clearly shows the incoherent adaptation of UDG on the target base, which is progressively pushed out of the pocket (see the steep increase of all plots in Fig.6f).

### 3.4 Is nucleotide-flipping possible in the nucleosome?

The literature about glycosylases identified unifying features of their mechanisms of action, as: (i) flipped-nucleotide structures accompanied by DNA bending; (ii) aromatic stacking and H-bonding interactions with specific features of the flipped nucleotide; (iii) enzyme side-chain intercalation into the DNA duplex to fill the empty slot left by the flipped nucleotide. Experiments allowed to identify a “search complex”, by which the glycosylase scans the DNA backbone, and an “interrogation complex”, by which the enzyme inspects the flipped-out nucleotide (see e.g. [42]). These features are observed in the static structures obtained by x-ray diffraction, nuclear magnetic resonance or cryo-microscopy studies, and indeed we found all such basic elements in the simulated molecular structures in the previous section. However, while static structures provide explanations for how enzymes recognize the specific features of damaged bases, they provide little insight into the dynamics of the process of recognition, and notably nucleotide-flipping, e.g.: how it is initiated; is it the result of a spontaneous, thermally-activated process or it is rather enzyme-induced by phosphodiester backbone interactions; are normal base-pairs also opened by the enzymes to ascertain their integrity, or are skipped after the search-complex?

As we just showed, rather strict steric conditions must be satisfied by the UDG/DNA interaction, in order for the excision chemistry to proceed, and the nucleosomal DNA presents a variety of structural arrangements that can probe this interaction. It was already demonstrated experimentally that base-excision efficiency is strongly dependent on the rotational position of the defects in the nucleosome [13, 14, 47]. By extrapolating from the results of the previous section for a typical DNA turn of about 11 base-pairs (Figure 7), only about 25% of the bases on average are in easily accessible positions (green), plus another 35% in difficult to access positions (yellow), the remaining 40% being permanently hidden (red). The largely reduced excision efficiency of UDG in the nucleosome compared to free DNA is clearly due to the complex arrangement of DNA around the histone core and, importantly, the role of histone tails. The enzyme can be outright interdicted access to the damage site, or can also be perturbed and deformed by its interaction with the nucleosome, to the extent that formation of the search- and interrogation-complex is severely hindered.

**Figure 7.**
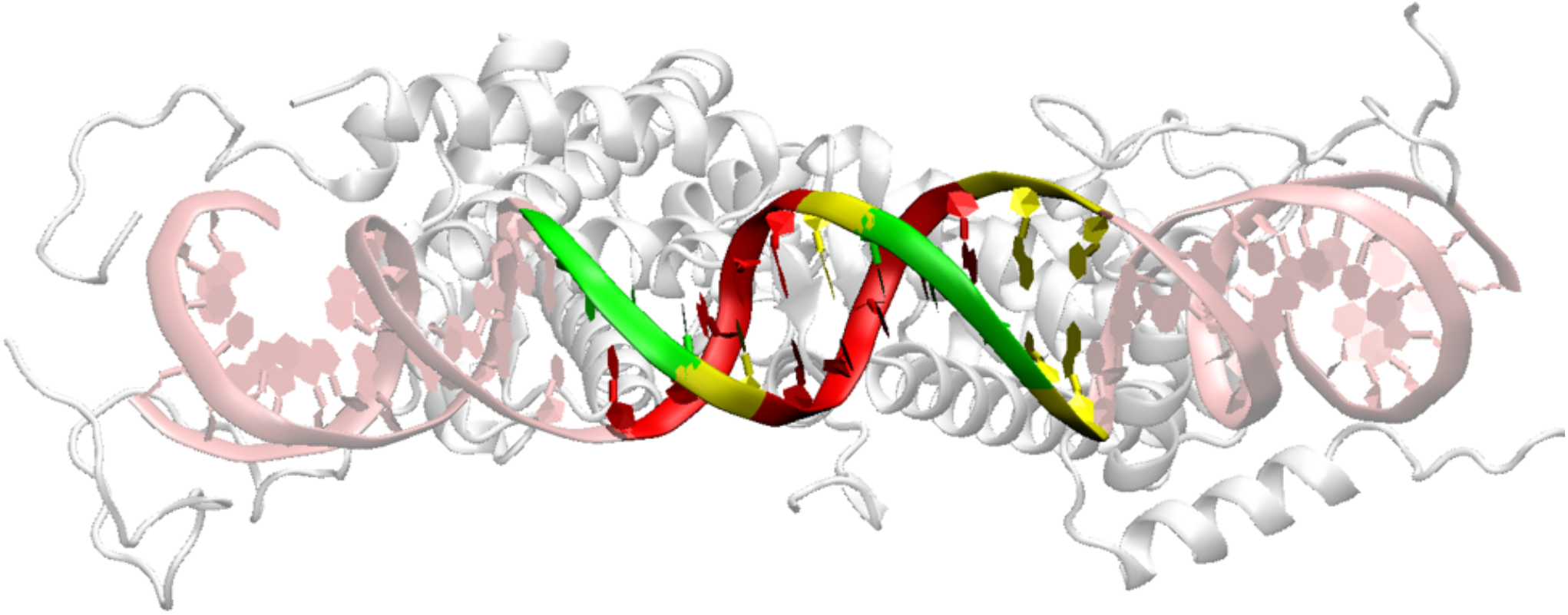
Schematic representation of about 1/3 of a nucleosome DNA loop (pink ribbons), with a half of histone core in the background (H3, H4, H2A, H2B as grey ribbons). Easily-accessible bases highlighted green, hardly-accessible yellow, hidden inaccessible red.

Such difficult (yellow) positions as the SHL–5, +6.5, or +1.5 described above, could be in theory available for docking a UDG. However, their constrained arrangement does not leave room for the target nucleotide to flip-out to form the interrogation complex, the entrance to the binding pocket being too narrow and blocked by one of the side loops (chiefly, His148). In these conditions, as shown by the above simulations, UDG could only passively adapt to a nucleotide already flipped-out by random thermal activation, while retaining a certain degree of base selectivity. Only “green” positions such as SHL-0, +3 or +4.5 allow for UDG to sit on the DNA, and form in rapid succession the search-complex, by fitting its Leu272 in the minor groove to help the flip-out of the target nucleotide, and the interrogation-complex, by accommodating the base of the flipped nucleotide deep inside the binding pocket.

To ascertain the ability of going dynamically from the search-to the interrogation-complex, we performed a few metadynamics simulations, by forcing extrahelical flipping of uracil at the ’easy’ positions SHL+3 or +4.5. In both cases, we started the simulations from the same docked UDG/DNA complex used as the initial configuration for the preceding 100-ns MD simulations, with the only modification of flipping back the target nucleotide (U34 or U46) to its normal intra-helical position. Upon application of the metadynamics bias potential, the target nucleotide explores the entire range of rotations by ±180° about the phosphate backbone, i.e. flipping both through the major and the minor groove. Clearly, the time trajectory in this case does not have a true kinematic meaning, however the most significative frames corresponding to local free-energy minima can be extracted, to analyze the molecular structures and reconstruct the most probable kinematics.

In both cases, the simple dihedral collective variable (CV) Y already used in our previous study [17,29] was not sufficient to obtain even an approximately correct interrogation-complex. For U34 at position SHL+3, the flipped base barely enters the binding pocket, however remaining at a distance from the active site. For U46 at position SHL+4.5, the access to the pocket is partially restrained by the proximity of His148, while Leu272 is not intercalating the minor groove due to the interference from the histone tail; as a result, the flipped nucleotide must make a torsion that keeps it from properly entering the binding pocket, albeit being flipped out by 180°.

To further accommodate the flipped nucleotide as close as possible to the interacting site, and make H-bonds with Asn204 and Phe158, it was necessary to add a second CV to the simulation. This was defined as the distance *d*_*CM*_ between the center of mass of the group Phe158-Asn204, and the center of the aromatic 6-ring of the uracil. While the dihedral CV takes care of the flipping, minimization of this distance should bring the flipped uracil close to the active site.

With such an arrangement it was possible for the case of U34 to obtain an optimal minimum free-energy configuration, in which the flipped uracil gets very close to the ideal position, with a total RMSD of <2 Å compared to the experimental **1EMH** (by best-fitting the position of the respective UDG). Figure 8a shows the free-energy map of the two CV, the dihedral Ψ varying between [-*π,π*] and the *d*_*CM*_ between [-1, 6] Å; the starting configuration is at Ψ = 0, *d*_*CM*_ = 1.5 Å; the absolute minimum is at Ψ = ±;,*π*, ∼*d*_*CM*_ 0.2 Å; note that the path via the major groove (Ψ *>* 0) is the most favorable, being mostly downhill in free-energy. The best interacting interrogation-complex (corresponding to the free-energy minimum indicated by a yellow ‘x’ in (a)) is shown in Fig.8b, where the simulated uracil is shown in blue and the experimental 1EMH is in red.

**Figure 8.**
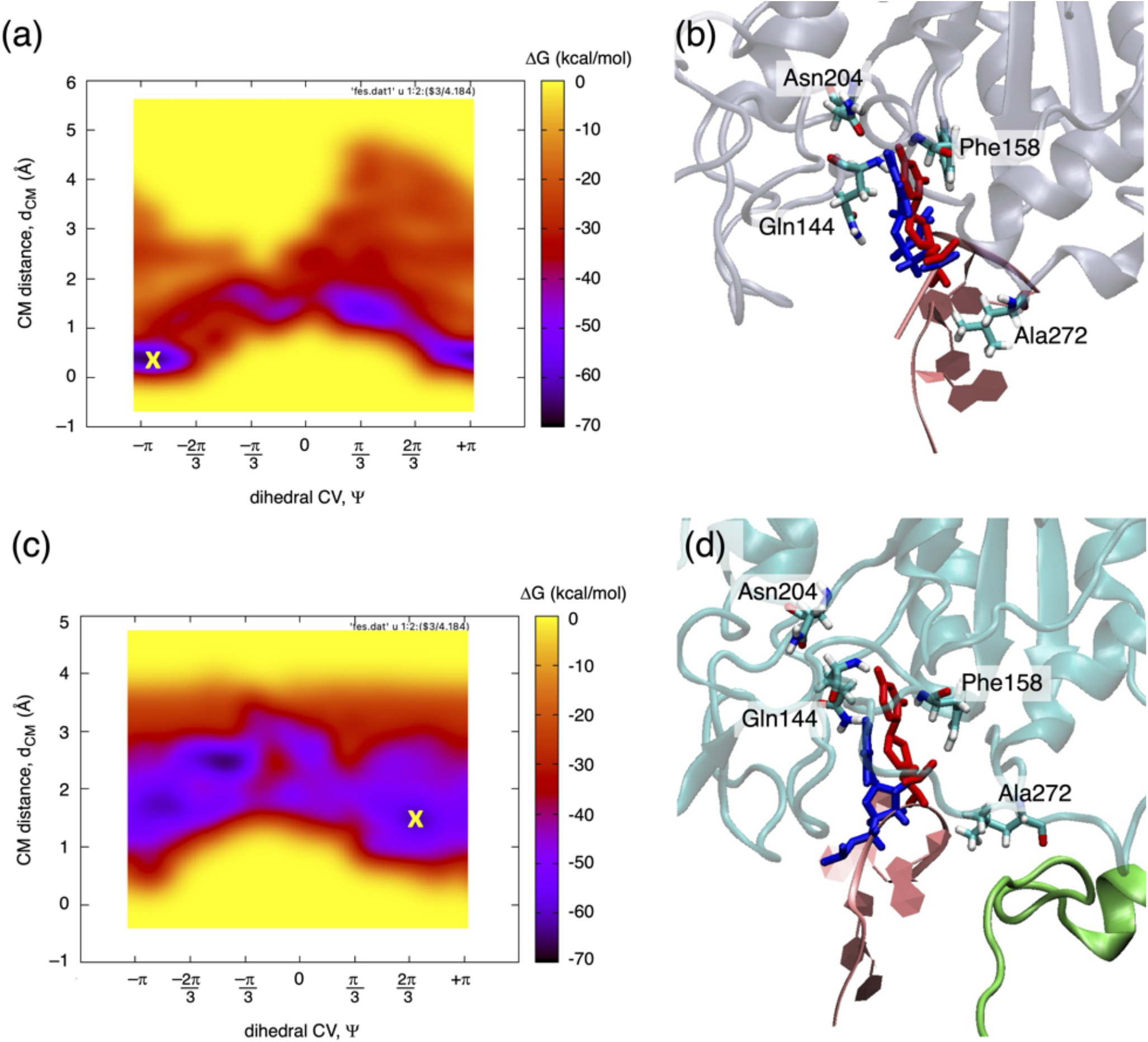
**(a**,**b)** Results of metadynamics simulation of the forced nucleotide flipping at position SHL+3, uracil U34. The left panel (a) shows the free-energy map in the plane of the two CV variables (Y, *d*_*CM*_). The right snapshot (b) corresponds to the free-energy minimum indicated by the yellow ‘x’ in (a); the simulated uracil is depicted in blue, the experimental 1EMH in red; the UDG is depicted in blue-grey ribbons in the background. **(c**,**d)** Same metadynamics results for uracil U46 at position SHL+4.5; the N-terminal of histone H2A is depicted in green ribbons in the foreground.

For the U46 uracil in position SHL+4.5, instead, even such a procedure could not find an interrogation-complex close enough to the experimental configuration. The free-energy map in Fig.8c displays several minima, none of which corresponding to values of *d*_*CM*_<1.5 Å; the absolute minimum is at Ψ ∼ -*π/*3, ∼ *d*_*CM*_ 2.5 Å. The best interrogation-complex, corresponding to the yellow ’x’ in (c), is shown in Fig.8d, has a total RMSD>20 Å: as it can be seen, the simulated uracil (blue) is quite off from the experimental position (red, obtained by best fitting the whole UDG in the two structures); the H2A histone tail (green) perturbs the correct intercalation of Leu272 in the DNA minor groove, this being likely the major steric obstacle to the correct positioning of U46 inside the binding pocket.

To further characterize the best configurations from metadynamics (those indicated by a yellow ’x’ in Fig.8), we calculated the DNA structural helical parameters with the help of the Curves+ program [48] (for the detailed geometrical definition of the helical parameters, see [49, 50]), with the results shown in Figure 9. The parameters are divided into intra-base-pair (open white symbols), that is relative to the deformation induced between the two bases in each pair, and inter-base-pair (grey shaded symbols), relative to the deformation between each two adjacent base-pairs.

**Figure 9.**
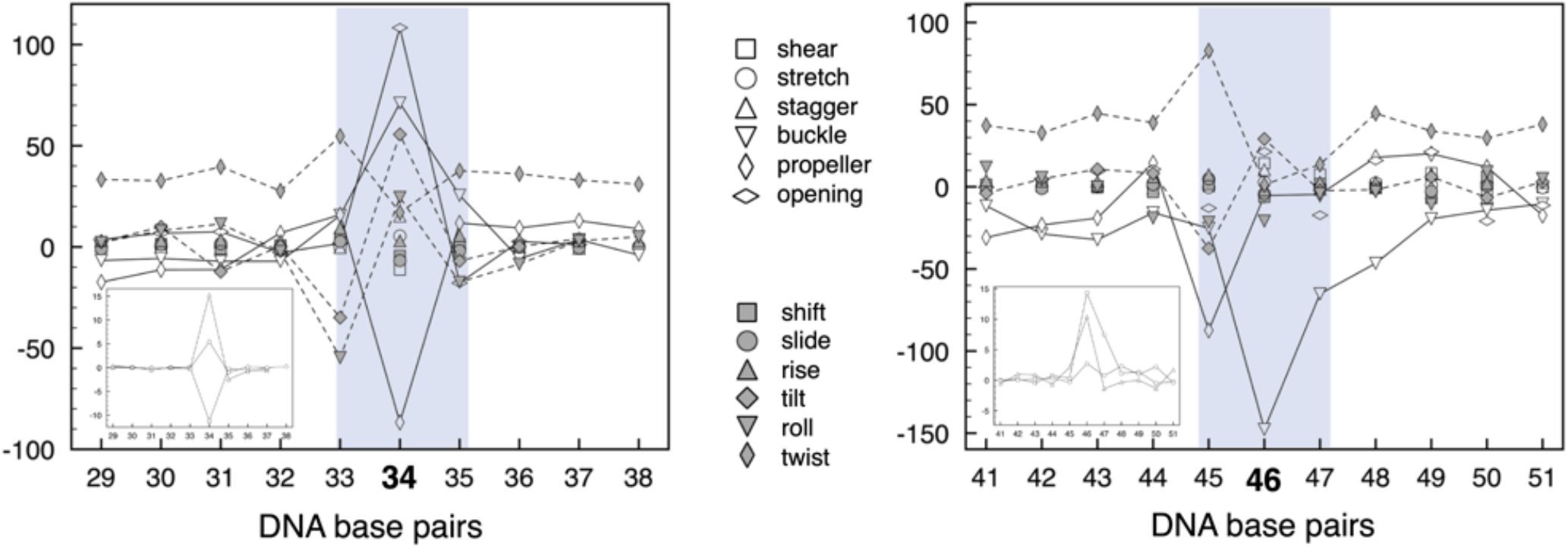
Structural helical parameters of DNA base pairs around the defect uracil, in the best configurations (the yellow ‘x’ in Fig.8) from the metadynamics simulation for the position SHL+3 (uracil U34) and position SHL+4.5 (uracil U46). Open symbols refer to inter-bp parameters, shaded symbols to intra-bp parameters. Vertical scale is Angstroms for translations, degrees for rotations. **Left, U34**. The inter-bp parameters buckle, opening, propeller are highlighted with a continuous line; the intra-bp parameters tilt, roll, twist with a dashed line. **Right, U46**. Buckle and propeller are highlighted with continuous line; tilt and twist with dashed line. The two insets display the inter-bp translational parameters with a better visible scale.

For U34 (left panel), a clear deviation from the average is observed just at the position 34 for the intra-bp parameters buckle, opening and propeller (highlighted by a continuous line): these are just the quantities defining the relative rotation between the two bases, about the local (x,y,z) reference frame. The inter-bp parameters tilt, roll and twist (highlighted by a dashed line) also describe the relative rotation between adjacent base-pairs: for tilt and roll (which also define the local DNA curvature, see Appendix in Ref. [50]) the two adjacent base-pairs 33 and 35 have a large negative fluctuation while the central base-pair including the flipped-out uracil has a large positive fluctuation; the opposite holds for the twist parameter, indicating large undertwisting at the bp 34, compensated by overtwisting at the adjacent bp 33 and 35. The inter-bp translational parameters (shown in the small inset) are also affected, but to a smaller extent, and the intra-bp translational parameters display even smaller deviations.

For U46 (right panel), for which the metadynamics could not find a fully correct interrogation-complex, it is seen a deviation from average for the buckle and propeller rotational degrees of freedom, however not fully localized at the bp 46 including the flipped-out nucleotide. Twist and tilt intra-bp parameters also display a similar behavior to U34, the deviations from average being less obvious in this case (for example, both the bp 46 and 47 exhibit undertwisting). Similar considerations as for the U34 hold for the both the intra-bp translational parameters (shown in the inset) and inter-bp parameters.

Overall, it appears that the rotational parameters are the ones carrying most of the information about the relative deformation induced by the flip-out, and enzyme binding processes, while the translational degrees of freedom are less affected; in all cases, the minor and major grooves are obviously widened at the flipped-out nucleotide site; both rotational and translational deformations are well localized to the flipped-out base pair in the well-fitting U34, while they tend to affect also neighboring base pairs in the imperfectly-fitting U46.

## 4 Discussion and Conclusions

In this work we studied the interaction of uracil-glycosylase repair protein with DNA uracil defects in the nucleosome by large-scale, all-atom molecular dynamics simulations. We focussed on the UDG enzyme that initiates the base-excision repair chain of DNA damage, being the very first actor in the identification of the damage site that intervenes to recognize and excise a wrongly incorporated uracil in the sequence.

The results of the simulations point out some interesting questions about the microscopic mechanisms of damage identification by UDG, and in general by glycosylases. First of all, it appears that only in a restricted number of situations an enzyme like UDG can find a correct arrangement on the DNA backbone, in order to form the search-complex and subsequently to move to the interrogation-complex. If we take the examples shown in section 3.3 as a representative sample of the different positions around the nucleosome, it turns out that only ∼25% of the bases around the nucleosome are readily accessible, while the rest is either inaccessible, or perturbed by the nucleosome structure, histone tails etc. Given that on average 75% of the genomic DNA is wrapped into nucleosomes, it would mean that less than 20% of the nucleosomal DNA is amenable to direct damage repair (to which one can add the 25% freely accessible, on average, in the linkers). Secondly, it appears that, even starting from a rather correct search-complex, however minor variances in the subsequent interrogation-complex induced by structural constraints can severely hamper the next chemical step leading to the excision-complex. In nearly all cases, however even when the complex is less accessible, UDG retains a substantial capability of discriminating between U and T. Third, and maybe more important, the ’hedgehog’-like nucleosome structure appears to make it difficult to imagine an enzyme like UDG freely sliding around the DNA backbone, while rapidly scouting tens of adjacent bases one after another, as it has been often assumed on the basis of studies on isolated DNA fragments [41, 42, 51]; enzyme sliding could possibly occur just on a very short range (1-2 bases next to the first one); instead, hopping between distant sites on a same nucleosome, or on a nearby one, should be the main search mechanism [52, 53]; the MD simulations of direct docking to a spontaneously-flipped and pre-bent nucleotide in section 3.3 also support this view (that is, closer to a “bend–then–bind”).

In addition to this, the metadynamics simulations of section 3.4 demonstrate that simple nucleotide flipping is not enough to obtain a correct interrogation-complex, but a second action besides flipping is necessary, to bring the flipped base well inside the binding pocket. In biophysical terms, this amounts to say that an extra force is necessary to deform and push the DNA strand, during or after the flipping. If we imagine an active role, or at least a participation of the enzyme in the deformation of DNA, in a kind of intermediate stage between the search- and the interrogation-complex, this could be identified with the “pinching” of the phosphate backbone, which in the case of UDG both experiments and our simulations attributed to the contact of serines 169 and 270. However, as we showed, this could work only in those cases in which UDG can readily find a perfect arrangement on the nucleosomal DNA. This is another factor that contributes to the largely reduced excision efficiency in the nucleosome, compared to free DNA [15, 16].

Given the severe mechanical (and entropic) constraints of the long DNA, twisted and wrapped around a tight succession of nucleosomes in the chromatin, it seems unlikely that damage may become accessible by purely thermally-activated events, such as nucleosome ’breathing’, sliding, or DNA reptation [54, 55]. These are fascinating hypothesis, but lacking experimental evidence, not the least because such events would lead to nucleosome disassembly in the chromatin [56]. In this context, nucleosome remodeling enzymes are far better candidates to lower the energy barriers that limit spontaneous nucleosome movements, by coupling the disruption of histone-DNA contacts to ATP hydrolysis (see e.g. [57, 58] and references therein). However, even active remodeling does not answer the main question, that is, *how* enzymes can efficiently identify DNA damage, if 3D diffusion is too slow, and 1D diffusion is restricted by the nucleosome structure and chromatin packing?

In our recent works [17, 18, 50, 59, 60], we proposed that localized mechanical deformation of DNA around the damaged site may be a proxy for signaling the presence of a defect. In the nucleosome, already several experiments [11, 61–65] demonstrated that the defect intrinsically increases local distortion, translation/rotational register shift, and partial opening of the DNA gyres. Moreover, large plastic (nonlinear) deformation around a DNA structural defect results from the constantly acting internal forces, of the order of hundreds of pN up to several nN, arising from chromatin mechanics and hydrodynamics [66–68], and chromatin interactions with the nuclear structural components [69, 70], notably (but not only) the peripheral lamins and nuclear membrane. The distortion induced by the redistribution of the accumulated elastic energy creates a long-ranged mechanical stress field that propagates along the DNA segment around the defect [50, 59]. These findings suggest that the complexity of the defect directly correlates with the degree of structural deformation and the amount of deformation energy injected: for example, in the present work we showed that DNA distortion in the case of a flipped-out nucleotide in the nucleosome is mostly concentrated in the rotational degrees of freedom; in the case of strand breaks [60] we found that stretch and shear translations carry most of the deformation; and in the case of DNA linkers [50] we showed that sharp kinking of DNA strands concentrates both the stress and excess elastic energy at the damage spot, while twist-bending redistributes the stress more smoothly about the zones of maximum DNA curvature. From a biological perspective, such structural alterations in damaged DNA, be it in the linker or around the nucleosome core, may serve as “markers”, exalting, as well as greatly accelerating, the initial damage recognition by damage-response enzymes, such as glycosylases, PARP, XPE/XPA, which could preferentially target regions of structural instability, rather than conducting an exhaustive search by diffusive mechanisms, randomly spanning back and forth along the (often inaccessible) chromatin DNA.

## Author contributions statement

All authors conceived the computer experiments. S.G. and F.C. conducted the computer simulations. F.C. drafted the manuscript. All authors analyzed the results and reviewed the manuscript.

## Acknowledgements

We acknowledge funding from the ANR Project “Dyprosome: dynamics of DNA repair proteins at nucleosomes” ANR-21-CE45-0032. We thank generous computing time allocation on the JEAN-ZAY supercomputer of IDRIS-CNRS in Orsay, and the ADASTRA supercomputer of CINES in Montpellier, under projects GENCI A0130712986 and A0150712986.

## Additional information

Partial trajectories and input files for the molecular dynamics simulations in this work are available at the repository FigShare, accession link: 10.6084/m9.figshare.28778600. Additional information available from the corresponding author.

The authors declare no competing interests.

## SUPPLEMENTARY MATERIALS

**Figure S1.**
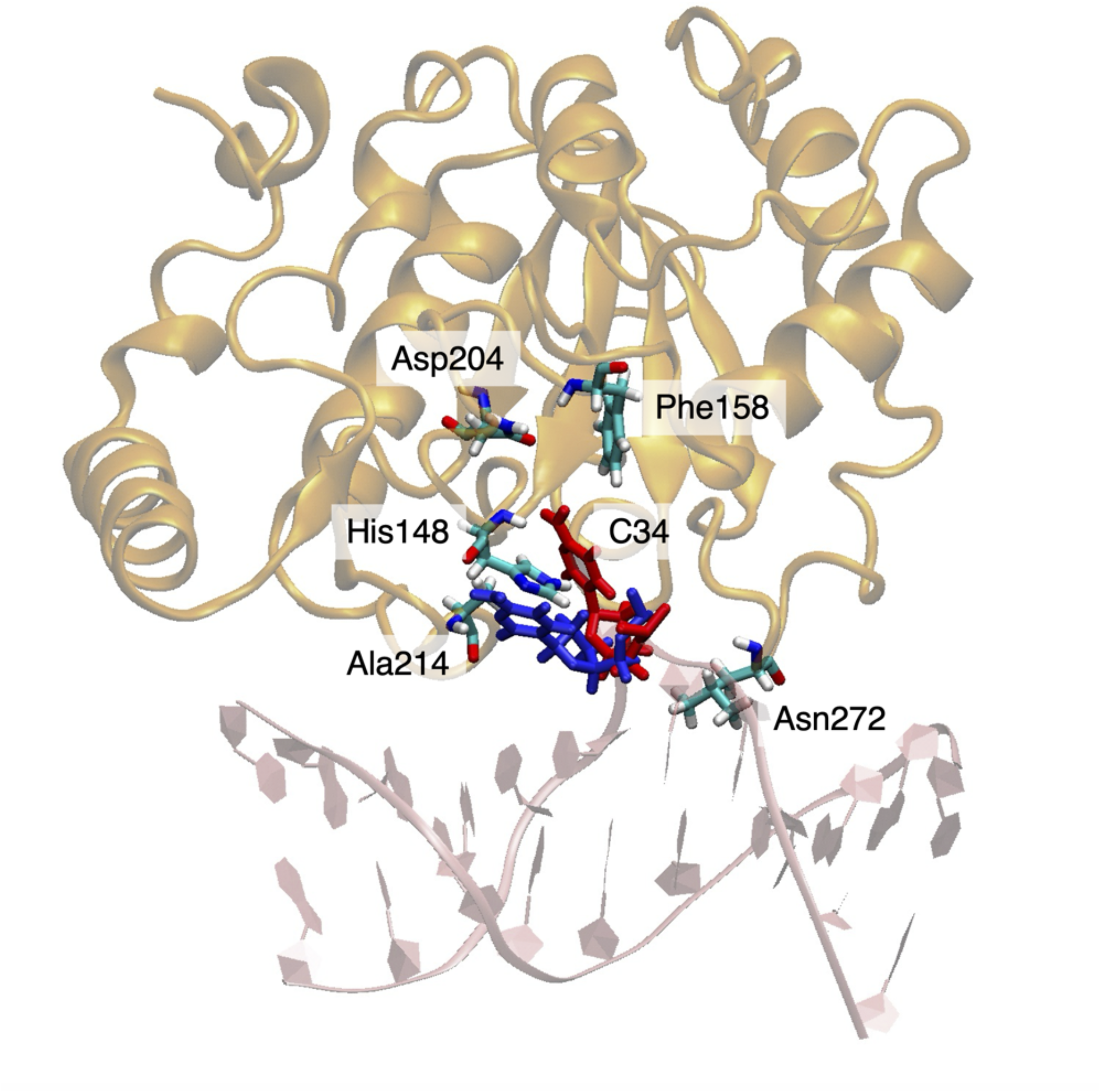
Position SHL+3, cytosine substitution for the flipped-out nucleotide. The initial position of C34 is shown in red sticks, and the position after 100-ns MD in blue sticks. UDG in orange ribbons, DNA in pink cartoon representation.

**Figure S2.**
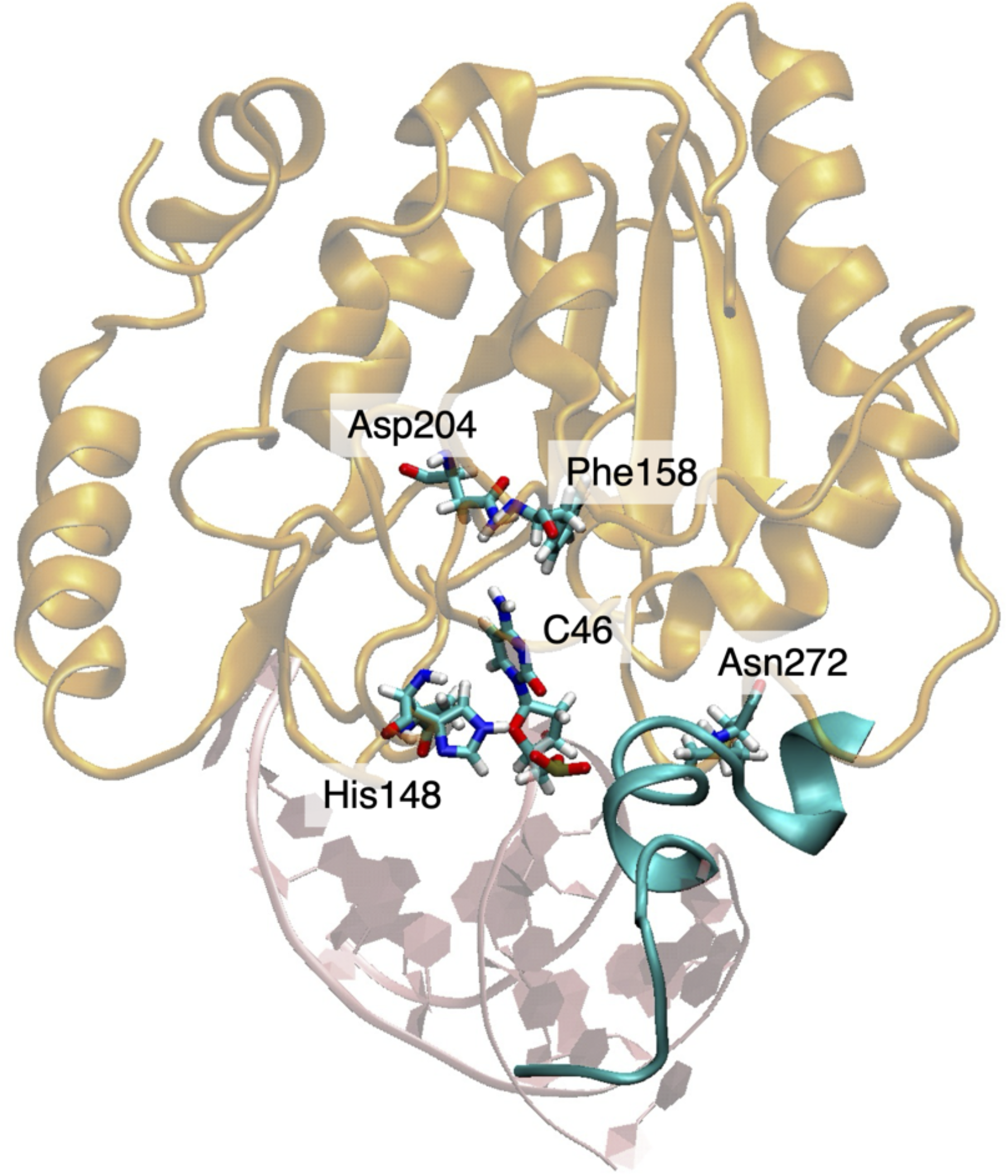
Position SHL+4.5, cytosine substitution for the flipped-out nucleotide. Initial configuration of C46 in the UDG binding pocket (orange ribbons); the N-terminal tail of histone H2A is also shown in cyan ribbons.

## References

1. Jacobs, A. L. & Schär, P. DNA glycosylases: in DNA repair and beyond. Chromosoma 121, 1–20 (2012).

2. Wallace, S. S., Murphy, D. L. & Sweasy, J. B. Base excision repair and cancer. Cancer Lett. 327, 73–89 (2012).

3. Guan, L. & Greenberg, M. M. Irreversible inhibition of DNA Polymerase b by an oxidized abasic lesion. J. Am. Chem. Soc. 132, 5004 (2010).

4. Jacobs, A. C., Kreller, C. R. & Greenberg, M. M. Long patch base excision repair compensates for DNA Polymerase b inactivation by the C40-oxidized abasic site. Biochemistry 50, 136–143 (2010).

5. Zhou, T. et al. Tyrosyl-DNA phosphodiesterase and the repair of 3’-phosphoglycolate-terminated DNA double-strand breaks. DNA Repair 8, 901–911 (2009).

6. Cao, C., Jiang, Y. L., Stivers, J. T. & Song, F. Dynamic opening of DNA during the enzymatic search for a damaged base. Nat Struct Mol Biol. 11, 1230–1236 (2004).

7. Yin, Y., Yang, L., Zheng, G. & Zhao, X. S. Dynamics of spontaneous flipping of a mismatched base in DNA duplex. Proc. Natl. Acad. Sci. (PNAS) 111, 8043–8048 (2014).

8. Sarangi, M. K. et al. Evidence for a bind-then-bend mechanism for architectural DNA binding protein yNhp6A. Nucl. Acids Res. 47, 2781–2791 (2019).

9. Panigrahi, A., Vemuri, H., Aggarwal, M., Pitta, K. & Krishnan, M. Sequence specificity, energetics and mechanism of mismatch recognition by DNA damage sensing protein Rad4/XPC. Nucl. Acids Res. 48, 2246–2257 (2020).

10. Yoshua, S. B. et al. Integration host factor bends and bridges DNA in a multiplicity of binding modes with varying specificity. Nucl. Acids Res. 49, 8684–8698 (2021).

11. Baral, S. et al. Evidence for intrinsic DNA dynamics and deformability in damage sensing by the Rad4/XPC nucleotide excision repair complex. Nucleic Acids Res. 53, gkae1290 (2025).

12. Cole, H. A., Tabor-Godwin, J. M. & Hayes, J. J. Uracil DNA glycosylase activity on nucleosomal DNA depends on rotational orientation of targets. J. Biol. Chem. 285, 2876–2885 (2010).

13. Ye, Y. et al. Enzymatic excision of uracil residues in nucleosomes depends on the local DNA structure and dynamics. Biochemistry 51, 6028–6038 (2012).

14. Rodriguez, Y. & Smerdon, M. J. The structural location of DNA lesions in nucleosome core particles determines accessibility by base excision repair enzymes. J. Biol. Chem. 288, 13863–13875 (2013).

15. Olmon, E. D. & Delaney, S. Differential ability of five DNA glycosylases to recognize and repair damage on nucleosomal DNA. ACS Chem. Biol. 12, 692–701 (2017).

16. Tarantino, M. E., Dow, B. J., Drohat, A. C. & Delaney, S. Nucleosomes and the three glycosylases: High, medium, and low levels of excision by the uracil DNA glycosylase superfamily. DNA Repair 72, 56–63 (2018).

17. Saravanan, V. et al. The ‘very moment’ when UDG recognizes a flipped-out uracil base in dsDNA. Sci. Reports 15, 7993 (2025).

18. Ghediri, S. et al. How repair proteins identify DNA damage in the nucleosome. Biomolecules X, ZZZ (2025).

19. Dominguez, C., Boelens, R. & Bonvin, A. M. HADDOCK: a protein-protein docking approach based on biochemical and/or biophysical information. J. Am. Chem. Soc. 125, 1731–1737, DOI: 10.1021/ja026939x (2006).

20. van Zundert, G. et al. The HADDOCK2.2 web server: User-friendly integrative modeling of biomolecular complexes. J. Mol. Biol. 428, 720–725, DOI: 10.1016/j.jmb.2015.09.014 (2015).

21. Yan, Y., Zhang, D., Zhou, P., Li, B. & Huang, S.-Y. HDOCK: a web server for protein–protein and protein–DNA/RNA docking based on a hybrid strategy. Nucl. Acids Res. 9, 988996, DOI: 10.1093/nar/gkx407 (2017).

22. Rodríguez-Lumbreras, L. A., Jiménez-García, B., Giménez-Santamarina, S. & Fernández-Recio, J. pyDockDNA: A new web server for energy-based protein-DNA docking and scoring. Front. Mol. Biosci. 9, 988996, DOI: 10.3389/fmolb.2022.988996 (2022).

23. Berendsen, H., van der Spoel, D. & van Drunen, R. GROMACS: A message-passing parallel molecular dynamics implementation. Comp. Phys. Comm. 91, 43–56, DOI: 10.1016/0010-4655(95)00042-E (1995).

24. Lindahl, E., Hess, B. & van der Spoel, D. GROMACS 3.0: a package for molecular simulation and trajectory analysis. J. Mol. Model. 7, 306–317, DOI: 10.1007/s008940100045 (2001).

25. Lindahl, E., Hess, B. & van der Spoel, D. ff14SB: Improving the accuracy of protein side chain and backbone parameters from ff99SB. J. Chem. Theor. Comp. 11, 3696–3713, DOI: 10.1021/acs.jctc.5b00255 (2015).

26. Ivani, I. et al. Parmbsc1: a refined force-field for DNA simulations. Nat. Meth. 13, 55–58, DOI: 10.1038/nmeth.3658 (2016).

27. Krissinel, E. & Henrick, K. Inference of macromolecular assemblies from crystalline state. J. Mol. Biol. 372, 774–797, DOI: 10.1016/j.jmb.2007.05.022 (2007).

28. Bonomi, M. et al. PLUMED: a portable plugin for free energy calculations with molecular dynamics. Comp. Phys. Comm. 180, 1961 (2009).

29. Song, K. et al. An improved reaction coordinate for nucleic acid base flipping studies. J. Chem. Theor. Comp. 5, 3105–3113 (2009).

30. Wang, H., Xiong, L. & Cramer, P. Structures and implications of TBP-nucleosome complexes. Proc. Natl. Ac. Sci. USA (PNAS) 118, e2108859118 (2021).

31. Duvaud, S. et al. Expasy, the Swiss Bioinformatics Resource Portal, as designed by its users. Nucl. Acids Res. 49, W216–W227, DOI: 10.1093/nar/gks225 (2021).

32. Lowary, P. & Widom, J. New DNA sequence rules for high affinity binding to histone octamer and sequence-directed nucleosome positioning. J. Mol. Biol. 276, 19–42, DOI: 10.1006/jmbi.1997.1494 (1998).

33. Stivers, J. T. Kinetic mechanism of damage site recognition and uracil flipping by Escherichia coli uracil DNA glycosylase. Biochemistry 38, 952–963 (1998).

34. Parikh, S. S., Putnam, C. D. & Tainer, J. Lessons learned from structural results on uracil-DNA glycosylase.. Mutat. Res. Repair 460, 183–199 (2000).

35. Parikh, S. et al. Uracil-DNA glycosylase-DNA substrate and product structures: conformational strain promotes catalytic efficiency by coupled stereoelectronic effects. Proceeding Natl. Acad. Sci. (PNAS) 97, 5083–5088 (2000).

36. Parikh, S. S. et al. Base excision repair initiation revealed by crystal structures and binding kinetics of human uracil-DNA glycosylase with DNA. EMBO J. 17, 5214–5226 (1998).

37. Pettersen, E. et al. UCSF-Chimera, a visualization system for exploratory research and analysis. J. Comput. Chem. 25, 1605–1612, DOI: 10.1002/jcc.20084 (2004).

38. Tye, B.-K., Nyman, P.-O., Lehman, I. R., Hochhauser, S. & Weiss, B. Transient accumulation of Okazaki fragments as a result of uracil incorporation into nascent DNA. Proceeding Natl. Acad. Sci. USA 74, 154–157 (1977).

39. Mol, C. et al. Crystal structure of human uracil-DNA glycosylase in complex with a protein inhibitor: protein mimicry of DNA. Cell 82, 701–708 (1995).

40. Earl, C. et al. A structurally conserved motif in gamma-herpesvirus uracil-DNA glycosylases elicits duplex nucleotide-flipping. Nucleic Acids Res. 46, 4286–4300 (2018).

41. Zharkov, D. O. & Grollman, A. P. The DNA trackwalkers: Principles of lesion search and recognition by DNA glycosylases. Mutat. Res. Mol. Mech. Mutagen. 57, 24–54 (2005).

42. Friedman, J. I. & Stivers, J. T. Detection of damaged DNA bases by DNA glycosylase enzymes. Biochemistry 49, 4957–4967 (2010).

43. da Rosa, G. et al. Sequence-dependent structural properties of B-DNA: what have we learned in 40 years? Biophys. Rev. 13, 995–1005, DOI: 10.1007/s12551-021-00893-8 (2021).

44. Assenza, S. & Pérez, R. Accurate sequence-dependent coarse-grained model for conformational and elastic properties of double-stranded DNA. J. Chem. Theory Comput. 18, 3239–3256, DOI: 10.1021/acs.jctc.2c00138 (2022).

45. Anderson, J. D. & Widom, J. Sequence and position-dependence of the equilibrium accessibility of nucleosomal DNA target sites. J. Mol. Biol. 296, 979–987 (2000).

46. Polach, K. J., Lowary, P. T. & Widom, J. Effects of core histone tail domains on the equilibrium constants for dynamic DNA site accessibility in nucleosomes. J. Mol. Biol. 298, 211–223, DOI: 10.1006/jmbi.2000.3644 (2000).

47. Odell, I. D., Wallace, S. S. & Pederson, D. S. Rules of engagement for base excision repair in chromatin. J. Cell Physiol. 228, 258–266 (2013).

48. Lavery, R., Moakher, M., Maddocks, J. H., Petkeviciute, D. & Zakrzewska, K. Conformational analysis of nucleic acids revisited: Curves+. Nucleic acids research 37, 5917–5929 (2009).

49. Olson, W. K. et al. A standard reference frame for the description of nucleic acid base-pair geometry. J. Mol. Biol. 313, 229–237, DOI: 10.1006/jmbi.2001.4987 (2001).

50. Cleri, F., Giordano, S. & Blossey, R. Nucleosome array deformation in chromatin is sustained by bending, twisting and kinking of linker DNA. J. Mol. Biol. 435, DOI: 10.1016/j.jmb.2023.168263 (2023).

51. Zharkov, D. O., Mechetin, G. V. & Nevinsky, G. A. Uracil-DNA glycosylase: Structural, thermodynamic and kinetic aspects of lesion search and recognition. Mutat. Res. /Fundamental Mol. Mech. Mutagen. 685, 11–20 (2010).

52. Porecha, R. H. & Stivers, J. T. Uracil-DNA glycosylase uses DNA hopping and short-range sliding to trap extrahelical uracils. Proc. Natl. Ac. Sci. USA (PNAS) 105, 10791–10796 (2008).

53. Hedglin, M. & O’Brien, P. J. Hopping enables a DNA repair glycosylase to search both strands and bypass a bound protein. ACS Chem. Biol. 5 (2010).

54. Widom, J. Equilibrium and dynamic nucleosome stability. Methods molecuar biology 119, 61–77 (1999).

55. Schiessel, H., Widom, J., Bruinsma, R. F. & Gelbart, W. M. Polymer reptation and nucleosome repositioning. Phys. Rev. Lett. 86, 4414–4417 (2001).

56. Rhee, H. S., Bataille, A. R., Zhang, L. & Pugh, B. F. Subnucleosomal structures and nucleosome asymmetry across a genome. Cell 159, 1377–1388, DOI: 10.1016/j.cell.2014.10.054 (2014).

57. Becker, P. B. Nucleosome sliding: facts and fiction. EMBO J. 21, 4749–4753, DOI: 10.1093/emboj/cdf486 (2002).

58. Ataian, Y. & Krebs, J. E. Five repair pathways in one context: chromatin modification during DNA repair. Biochem. Cell Biol. 84, 490–504 (2006).

59. Cleri, F., Landuzzi, F. & Blossey, R. Mechanical evolution of DNA double-strand breaks in the nucleosome. PLOS Comp. Biol. 14, e1006224, DOI: 10.1371/journal.pcbi.1006224 (2018).

60. Sarma, P. A., Abbadie, C. & Cleri, F. Cooperative dynamics of PARP1 zinc-finger domains in the detection of DNA single-strand breaks. Sci. Reports 14, 23257 (2024).

61. Sultanov, D. et al. Unfolding of core nucleosomes by PARP1 revealed by spFRET microscopy. AIMS genetics 4, 021–031 (2017).

62. Matsumoto, S. et al. DNA damage detection in nucleosomes involves DNA register shifting. Nature 571, 79–84, DOI: 10.1038/s41586-019-1259-3 (2019).

63. Maluchenko, N. V. et al. Mechanisms of nucleosome reorganization by PARP1. Int. J. Mol. Sci. 22, 12127 (2021).

64. Weaver, T. M. et al. Structural basis for APE1 processing DNA damage in the nucleosome. Nat. Commun. 13, 5390, DOI: 10.1038/s41467-022-33057-7 (2022).

65. Zheng, L., Tsai, B. & Gao, N. Structural and mechanistic insights into the DNA glycosylase AAG-mediated base excision in nucleosome. Cell Discov. 9, 62, DOI: 10.1038/s41421-023-00560-0 (2023).

66. Blossey, R. & Schiessel, H. The dynamics of the nucleosome: thermal effects, external forces and ATP. FEBS J. 278, 3619–3632, DOI: 10.1111/j.1742-4658.2011.08283.x (2011).

67. Bruinsma, R., Grosberg, A. Y., Rabin, Y. & Zidovska, A. Chromatin hydrodynamics. Biophys. J. 106, 1871–1881 (2014).

68. Dupont, S. & Wickström, S. A. Mechanical regulation of chromatin and transcription. Nat. Rev. Genet. 23, 624–643 (2022).

69. Amar, K., Wei, F., Chen, J. & Wang, N. Effects of forces on chromatin. APL Bioeng. 5, 041503 (2021).

70. Hsia, C.-R., Melters, D. P. & Dalal, Y. The force is strong with this epigenome: Chromatin structure and mechanobiology. J. Mol. Biol. 435, 168019 (2023).

